# Copy number variations shape the structural diversity of Arabidopsis metabolic gene clusters and are associated with the climatic gradient

**DOI:** 10.1101/2022.09.05.506152

**Authors:** Malgorzata Marszalek-Zenczak, Anastasiia Satyr, Pawel Wojciechowski, Michal Zenczak, Paula Sobieszczanska, Krzysztof Brzezinski, Tetiana Iefimenko, Marek Figlerowicz, Agnieszka Zmienko

**Affiliations:** Institute of Bioorganic Chemistry, Polish Academy of Sciences, Poland; Institute of Computing Science, Faculty of Computing and Telecommunications, Poznan University of Technology, Poznan, Poland; National University of Kyiv-Mohyla Academy, Kyiv, Ukraine

**Keywords:** copy number variation, biosynthetic gene cluster, secondary metabolism, coexpression, oxidosqualene cyclase, GWAS

## Abstract

Metabolic gene clusters (MGCs) encode at least three different enzymes for a common biosynthetic pathway. Comparative genome analyses highlighted the role of duplications, deletions and rearrangements in MGC formation. We hypothesized that these mechanisms also contribute to MGC intraspecies diversity and play a role in adaptation. We assessed copy number variations (CNVs) of four *Arabidopsis thaliana* MGCs in a population of 1,152 accessions, with experimental and bioinformatic approaches. The MGC diversity was lowest in marneral gene cluster (one private deletion CNV) and highest in the arabidiol/baruol gene cluster where 811 accessions had gene gains or losses, however, there were no presence/absence variations of the entire clusters. We found that the compact version of thalianol gene cluster was predominant in *A. thaliana* and more conserved than the noncontiguogus version. In arabidiol/baruol cluster we found a large insertion in 35% of analyzed accessions, that contained duplications of the reference genes *CYP705A2* and *BARS1*. The *BARS1* paralog, which we named *BARS*2, encoded a novel oxidosqualene synthase. Unexpectedly, in accessions with the insertion, the arabidiol/baruol gene cluster was expressed not only in roots but also in leaves. Additionally, they presented different root growth dynamics and were associated with warmer climates compared to the reference-like accessions. We also found that paired genes encoding terpene synthases and cytochrome P450 oxidases had higher copy number variability compared to non-paired ones. Our study highlights the importance of intraspecies variation and nonreference genomes for dissecting secondary metabolite biosynthesis pathways and understanding their role in adaptation and evolution.

## Introduction

Plants are able to produce a variety of low molecular weight organic compounds, which enhance their ability to compete and survive in nature. Secondary metabolites are not essential for plant growth and development. However, they are often multifunctional and may act both as plant growth regulators and be engaged in the primary metabolism or plant protection (Isah, 2019; Erb and Kliebenstein, 2020). The ability to produce particular types of compounds is usually restricted to individual species or genera. Therefore, these compounds are enormously diverse and have a wide range of biological activity. In plants, genes involved in a common metabolic pathway are typically dispersed across the genome. In contrast, functionally related genes that encode the enzymes involved in specialized metabolite biosynthesis in bacteria and fungi are frequently coexpressed and organized in the so-called operons (Boycheva et al., 2014; Nützmann et al., 2018). Similar gene organization units named biosynthetic gene clusters or metabolic gene clusters (MGCs) have been recently found in numerous plant species. MGCs are classically defined as a group of three or more genes, which i) encode a minimum of three different types of biosynthetic enzymes ii) are involved in the consecutive steps of a specific metabolic pathway and iii) are localized in adjacent positions in the genome or are interspersed by a limited number of intervening (i.e., not functionally related) genes (Nützmann and Osbourn, 2014; Kautsar et al., 2017). A typical MGC contains a “signature” enzyme gene involved in the major (usually first) step of a biosynthetic pathway. In this step, the metabolite scaffold is generated that determines the class of the pathway products (e.g., terpenes or alkaloids). This scaffold is further modified by “tailoring” enzymes encoded by other clustered genes, e.g. cytochrome P450 oxidases (CYPs), acyltransferases or alcohol dehydrogenases. The contribution of other enzymes encoded by peripheral genes (i.e. located outside the MGC) and the connection network between different metabolite biosynthesis pathways may result in additional diversification of the biosynthetic products (Huang et al., 2019). Currently, there are over 30 known MGCs in plants from various phylogenetic clades and new ones are being discovered. Their sizes range from 35 kb to several hundred kb. However, clusters of functionally related nonhomologous genes are still considered unusual in plant genomes.

In *Arabidopsis thaliana* (hereafter Arabidopsis), four MGCs have been discovered so far (**Supplemental Table S1**). They are involved in the metabolism of specialized triterpenes: thalianol, marneral, tirucalladienol, arabidiol and baruol. Triterpenes constitute a large and diverse group of natural compounds derived from 2,3-oxidosqualene cyclization, in a reaction catalyzed by oxidosqualene cyclases (OSCs) (Thimmappa et al., 2014). Out of 13 OSC genes known in the Arabidopsis genome, five (*THAS1, MRN1, PEN3, PEN1, BARS1*) are located within MGCs and encode their “signature” enzymes (Field and Osbourn, 2008; Field et al.; 2011; Boutanaev et al., 2015). The thalianol gene cluster contains five members, involved in the thalianol production and its conversion to another triterpene, thalianin (Fazio et al., 2004; Field and Osbourn, 2008; Huang et al., 2019). In the reference genome, this MGC is ∼45 kb in size. Thalianol synthase gene *THAS1, CYP708A2, CYP705A5* and *AT5G47980* (BAHD acyltransferase) genes are tightly clustered together, with only one noncoding transcribed locus (*AT5G07035*) between them. The fifth member, acyltransferase *AT5G47950*, is separated from the rest of the cluster by *RABA4C* and *AT5G47970* intervening genes. The marneral gene cluster is ∼35 kb in size and is the most compact plant MGC described to date. It is made up of three members: marneral synthase gene *MRN1*, marneral oxidase gene *CYP71A16* and *CYP705A12*, which function is unknown (Xiong et al., 2006; Field et al., 2011; Go et al., 2012). Additonally, there are three noncoding transcribed loci (*AT5G00580, AT5G06325* and *AT5G06335*) located between *CYP701A16* and *MRN1*. The tirucalladienol gene cluster is ∼47 kb in size and includes five members: tirucalla-7,24-dien-3β-ol synthase gene *PEN3*, an uncharacterized acyltransferase gene *SCPL1*, which was identified based on its co-expression with *PEN3, CYP716A1*, which is involved in the hydroxylation of tirucalla-7,24-dien-3β-ol, as well as *AT5G36130* and *CYP716A2* (Morlacchi et al., 2009; Boutanaev et al., 2015; Wisecaver et al., 2017). The contiguity of this MGC is interrupted by four intervening genes (*CCB3, AT5G36125, HCF109* and *AT5G36160*) and the noncoding locus *AT5G05325*. The arabidiol/baruol gene cluster is most complex and has estimated size of 83 kb. It encompasses two closely located OSCs, *PEN1* and *BARS1*, sharing 91% similarity at the amino acid level. *BARS1* encodes a multifunctional cyclase, which produces baruol as its main product (Lodeiro et al., 2007). *PEN1* encodes arabidiol synthase and is adjacent to *CYP705A1*, which is involved in arabidiol degradation upon jasmonic acid treatment (Xiang et al., 2006; Castillo et al., 2013; Sohrabi et al., 2015). The role of the remaining genes in arabidiol/baruol gene cluster (*CYP702A2, CYP702A3, CYP705A2, CYP705A3, CYP705A4, CYP702A5, CYP702A6* as well as acyltransferases *AT4G15390* and *BIA1*) has not been determined, however, they displayed coexpression with either *PEN1* or *BARS1* (Wada et al., 2012; Wisecaver et al., 2017). There are few intervening loci in arabidiol/baruol gene cluster, including a protein-coding gene *CSLB06*, two pseudogenes *CYP702A4P* and *CYP702A7P* and one novel transcribed region *AT4G06325*.

Plant MGCs are thought to have arisen by duplication and subsequent neo- or sub-functionalization of genes involved in the primary metabolism, which might have been followed by the recruitment of additional genes to the newly forming biosynthetic pathway (Nützmann and Osbourn, 2014). MGCs are frequently located within dynamic chromosomal regions, e.g., sub-telomeric, centromeric or rich in transposable elements (TEs), where the possibility of bringing together the beneficial sets of genes by structural rearrangements may be higher than in the rest of the genome, thus promoting MGC formation (Field et al., 2011). However, the same factors may also contribute to further genetic modifications and alteration of plant metabolic profile, thus making such MGCs “evolutionary hotspots”. To verify this scenario, we evaluated the intraspecific diversity of Arabidopsis MGCs and examined whether this diversity is associated with the trait variation. Here, we present a detailed picture of MGC copy number variations (CNVs), describe the discovery of novel, nonreference genes in arabidiol/baruol gene cluster and reveal the links between the variation in MGC structure and plant adaptation to different natural environments.

## Results

### MGCs differ in levels of copy number polymorphism

We started our analysis from aligning each MGC with the common CNVs in Arabidopsis genome, which were identified previously (Zmienko et al., 2020). As expected, each MGC had a substantial overlap with the variable regions: 100% for the thalianol gene cluster, 79.6% for the tirucalladienol gene cluster, 53.1% for the arabidiol/baruol gene cluster, and 52.8% for the marneral gene cluster (**Figure 1A**). However, the potential impact of CNVs on the clustered genes differed among the MGCs (**Supplemental Figure S1; Supplemental Table S2**). In thalianol gene cluster, most CNVs were grouped in the region spanning *AT5G47980, CYP705A5, CYP708A2* and *THAS1*, while *AT5G47950* was covered only by the largest variant CNV_18592 (241 kb in size), which encompassed the entire cluster. In the arabidiol/baruol gene cluster, the CNVs (0.6 kb to 21 kb in size) were grouped in three distinct regions, separated by invariable segments. The first variable region overlapped with *CYP702A2* and *CYP702A3*. The second variable region overlapped with *CYP705A2, CYP705A3* and *BARS1*. The CNVs in the third variable region were mostly intergenic and overlapped only two genes, *CYP702A5* and *CYP702A6. CYP705A1, PEN1, CYP705A4, AT4G15390* and *BIA1* were not covered by any common CNV. In the tirucalladienol gene cluster, the CNVs accumulated in the 5’ part of the cluster and none of them overlapped with *SCPL1*.

**Figure 1.**
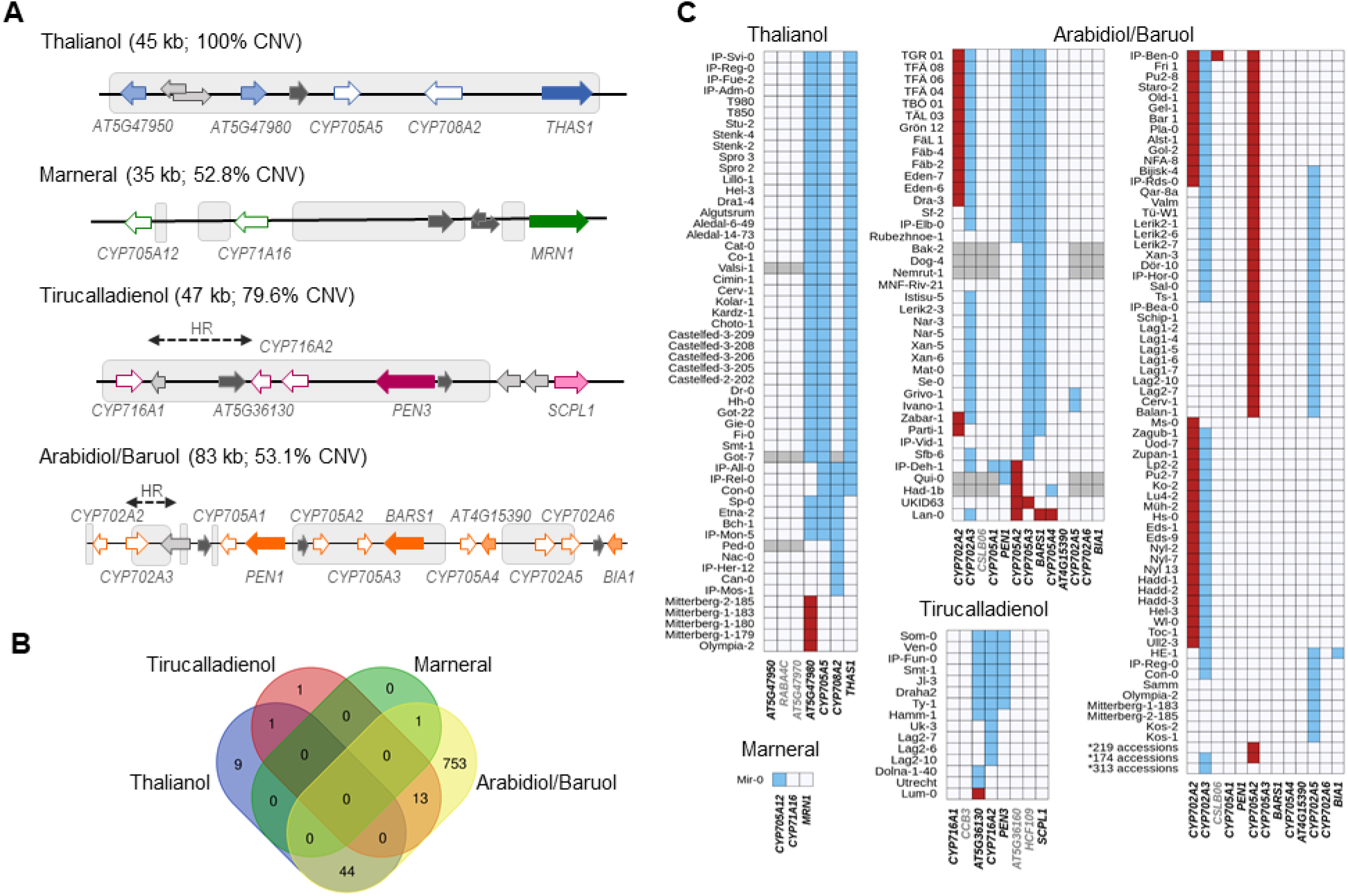
Copy number variation of Arabidopsis metabolic gene clusters. A) MGC overlap with CNV regions. Colored arrows with white filling denote CYPs. Arrows with dark color filling denote OSCs. Arrows with light color filling denote other types of MGC genes. Intervening genes are in light grey. Noncoding genes are in dark grey. Grey boxes indicate overlap with CNV regions. HR – hotspot of rearrangements; B) Number and overlap among the accessions with detected gene copy numbers in each of four MGCs; C) Patterns of gene copy number variation in each MGC. Red – gain; blue – loss, grey – no assignment. Names of the genes considered as MGC members are in black; names of the intervening genes are in grey. Source data for histograms are in Supplemental Table S6.

Noteworthy, upstream of the tirucalladienol gene cluster, a region genetically divergent from the surrounding genomic segments, called a hotspot of rearrangements, was previously described (Jiao and Schneeberger, 2020). Smaller hotspots of rearrangements were also found between *CYP716A1* and *AT5G36130* in the same MGC as well as in one variable segment of arabidiol/baruol gene cluster. Their occurence stayed in agreement with the high number of CNVs in these genomic regions. The CNV arrangement in marneral gene cluster was strikingly different from any other MGC, in that all variants were intergenic and did not overlap with the marneral cluster genes.

Nonuniform distribution of CNVs in MGC regions indicated that the triterpene biosynthetic pathways encoded by these MGCs in different accessions may be non-identical. To evaluate this possibility, we retrieved copy number data for 31 genes (clustered and intervening genes in all MGCs), each from 1,056 accessions (RD dataset; **Supplemental Table S3**) and supplemented them with multiplex ligation-dependent amplification assays for 232 accessions (MLPA dataset; **Supplemental Table S4**) and droplet digital PCR-based genotyping assays for 20 accessions (ddPCR dataset; **Supplemental Table S5**). We defined the thresholds for detecting duplications and deletions for each data type. Next, we assigned the copy number status of each gene in each accession (“REF”, “LOSS” or “GAIN”) by combining all three datasets (**Supplemental Table S6**). Out of the genotypes assigned with two or three approaches, 98.8% were fully concordant and most of the remaining discrepancies could be resolved manually (**Supplemental Figures S2-S4; Supplemental Table S7**). The combined genotyping data for 1,152 accessions were further used to asses and compare MGC variation at the gene level.

Only 28.6% of the assayed accessions had no gene gains or losses in any MGC (**Figure 1B**). This included 65% of accessions from the German genetic group and 39% of accessions from the Central Europe group. Contrarily, the vast majority (at least 90%) of accessions from groups known to be genetically distant from the reference genome (North Sweden, Spain, Italy-Balkan-Caucasus, and Relict groups) displayed gene CNV in at least one MGC. We note that the real number of invariable accessions could be even lower since for 96 accessions some MGC genes were not genotyped.

Altogether, 19 genes were affected: four in the thalianol cluster, one in the marneral cluster, three in the tirucalladienol cluster and 11 in the arabidiol/baruol cluster (**Figure 1C**). The latter was also most variable in terms of the number of accessions carrying CNVs and the diversity of CNV patterns. For two genes we detected only copy gains, for 11 – only losses, while six genes were multiallelic (with both gains and losses). As expected, these genes resided in the previously defined variable regions. Remarkably, we did not observe complete loss or gain of entire MGC in any accession. In the next step, we inspected in more detail the level of diversity of each MGC.

### The compact version of thalianol gene cluster is predominant and more conserved than the reference-like noncontiguous version

Survey with a combination of RD, MLPA and ddPCR approaches revealed 54 accessions with the copy number changes in the thalianol gene cluster, which followed five distinct patterns and *AT5G47950* was the only invariant gene in all accessions (**Figure 2A**). The most common (variant A) was the deletion of a region encompassing *AT5G47980* and *CYP705A5*, combined with the deletion of *THAS1*. We detected this variant in 37 accessions from six countries: Sweden (13), Italy (8), Germany (6), Spain (5), Bulgaria (3) and Portugal (2). We also confirmed the existence of two previously reported rare variants (Liu et al., 2020a). One of them (variant B) was a large deletion spanning *AT5G47980, CYP705A5* and *CYP708A2*. We found this variant in two accessions from Germany (Bch-1, Sp-0), one from Italy (Etna-2) and one from Spain (IP-Mon-5). The other one (variant C) was a deletion of a single gene, *CYP708A2*, which we found in five accessions, mainly Relicts, originating from Spain (Can-0, Ped-0, IP-Her-12 and Nac-0) and Portugal (IP-Mos-1). We also found a new type of deletion (variant D) in two Spanish Relicts (IP-Rel-0 and Con-0) and one non-Relict (IP-All-0). The deletion spanned *CYP705A5, CYP708A2* and *THAS1* **(Supplemental Figure S5**). The last variant (variant E) was a duplication of acyltransferase gene *AT5G47980*, which was found in four accessions from Italy (Mitterberg-1-179, Mitterberg-1-180, Mitterberg-1-183, Mitterberg-2-185) and one from Greece (Olympia-2). The presence of a tandem duplication ∼3kb in size in Mitterberg-2-185 was confirmed by sequence analysis of its *de novo* genomic assembly **(Supplemental Figure S6**). The duplication spanned entire *AT5G47980* and its flanks (0.5 kb upstream and 0.7 kb downstream) and differed from its copy only by two mismatches and a 1-bp gap. The predicted protein products of both gene copies were identical and shorter than the reference acyltransferase (404 aa versus 443 aa), but they possessed complete transferase domains (pfam02458).

**Figure 2.**
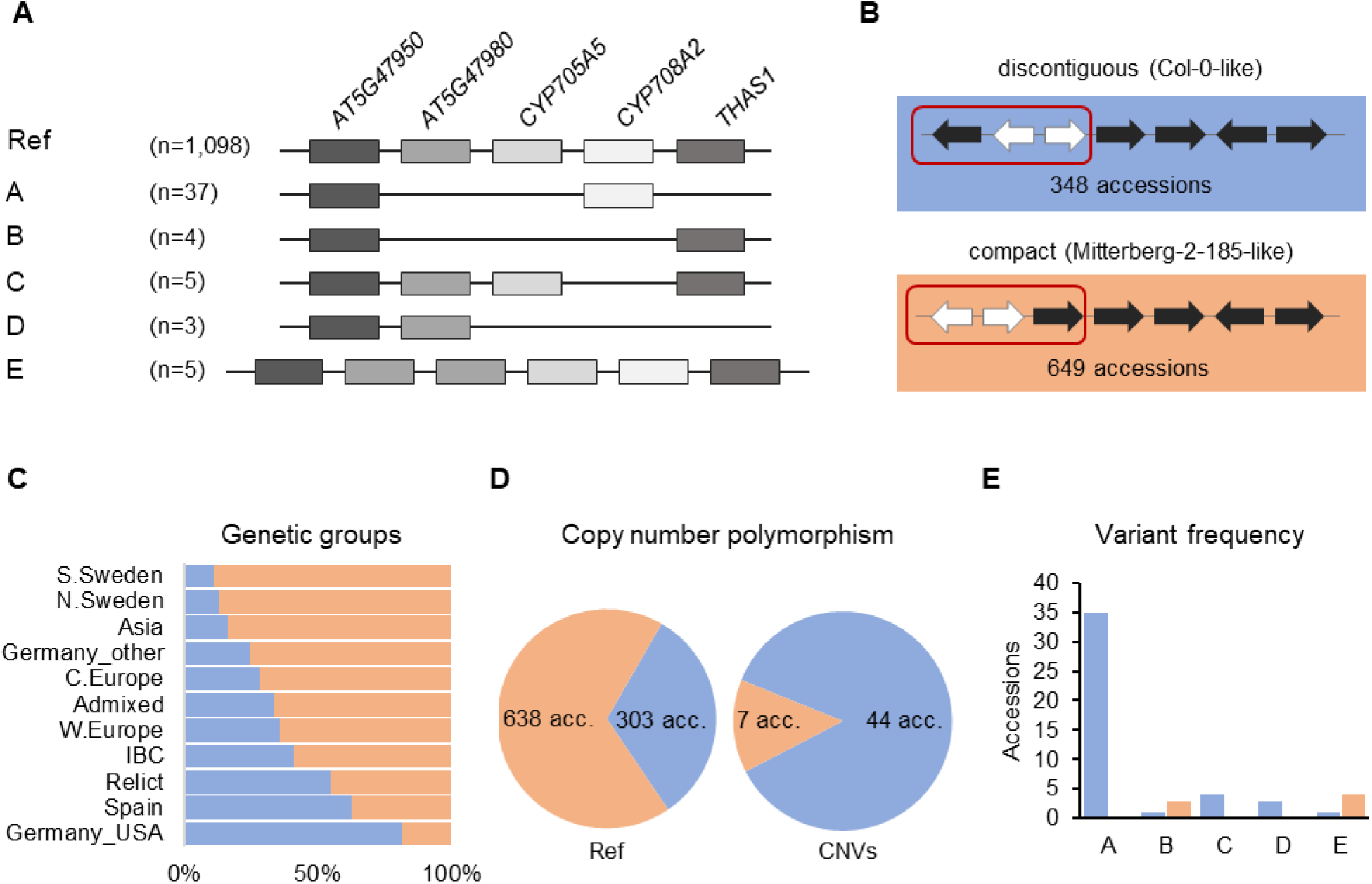
Structural variation of thalianol gene cluster. A) Five types of CNVs that change the number of thalianol cluster genes. The position of intervening genes is ignored and they are not shown. Gene orientation is disregarded. B) Two versions of thalianol gene cluster organization. Clustered genes are in black; interfering genes are in white. C) The frequency of the two thalianol gene cluster versions (discontiguous and compact) among the genetic groups. D) Rate of copy number polymorphism within discontiguous and compact clusters. E) Frequency of variants presented in (A) among the accessions with different cluster organizations. The number of presented accessions in panels is 1,152 for (A) – genotyping, 997 for (B,C) – inversion detection and 992 for (D,E) – the intersection of the above.

In Mitterberg-2-185 assembly we also detected a chromosomal inversion (with respect to the reference genome orientation), spanning *AT5G47950* and two intervening genes, *RABA4C* and *AT5G47970*. It resulted in a more compact cluster organization compared to the reference (**Figure 2B**). Similar inversions were previously detected in 17 other accessions (out of 22 analyzed), which indicated that the compact version of thalianol gene cluster might be predominant in Arabidopsis (Liu et al., 2020a). To verify this possibility, we set up a bioinformatic pipeline for detecting genomic inversions based on paired-end genomic reads analysis, in 997 accessions. We correctly detected inversions in 12 out of 15 previously analyzed accessions, which indicated good sensitivity of our method. Altogether, we found inversions, 12.8 kb to 15.4 kb in size, spanning *AT5G47950, RBAA4C* and A*T5G47970* genes, in 649 accessions (65%), which fully confirmed our predictions (**Supplemental Table S8**). The compact version of thalianol gene cluster was dominant in South and North Sweden genetic groups as well as Asia group (83.6% to 88.9%), while the discontiguous version was mainly observed among U.S.A accessions and was also slightly more abundant in Spain genetic group (**Figure 2C**). There was similar frequency of discontiguous and compact versions among the Relicts (12 and 10 accessions, respectively). Interestingly, the CNV frequency substantially differed between the accessions with different cluster organization (**Figure 2D, E**). The compact cluster was more conserved - copy number changes (variants B and E) affected only 1.1% of accessions in this group. The remaining variants including deletions spanning *THAS1* signature gene, were found exclusively among the accessions with the reference-like cluster type. Altogether, 12.7% of accessions with the discontiguous cluster were affected by CNVs.

### Marneral and tirucalladienol gene clusters display little structural variation

Analysis of RD and MLPA data confirmed exceptionally low variability of marneral cluster genes. One private variant, which we detected in Mir-0 and confirmed by Sanger sequencing, was 1.2 kb in size and spanned the first exon of *CYP705A12* gene, which resulted in the truncation of its predicted protein product **(Supplemental Figure S7**). Apart from that, we did not detect any common gene duplications or deletions within this MGC. Likewise, we observed low variation of tirucalladienol gene cluster. In 15 accessions (1.4%), deletions or duplications occurred in the region spanning *AT5G36130, CYP716A2* and *PEN3* genes and affected one, two or all of them. Differences between the countries indicated these structural variants weree of local origin (**Supplemental Figure 8**). Sequence analysis of *de novo* genomic assemblies for Ty-1 and Dolna-1-40 confirmed the predicted deletion patterns in these accessions. It should be noted that, according to a recent study, *AT5G36130* and *CYP716A2* gene models are misannotated and they jointly encode a single protein of the CYP716A subfamily with cytochrome oxidase activity (Yasumoto et al., 2016) (**Supplemental Figure S9**). Therefore, a full-length gene was absent from all 15 accessions with CNVs in tirucalladienol gene cluster (**Figure 1C**).

### Intraspecies variation of arabidiol/baruol gene cluster reveals a novel OSC gene

The arabidiol/baruol gene cluster was the most heterogeneous of all MGCs. Consistent with the segmental CNV coverage, there were apparent differences in the variation frequency between the genes. At the cluster’s 5’end, *CYP702A2* was duplicated in 50 accessions and *CYP702A3* was deleted in 564 accessions, including about 70% of all analyzed accessions from Sweden and Spain. On the contrary, genes located at the 3’ end of the cluster showed little variation. There were *CYP702A5* deletions in 35 accessions, *CYP705A4* deletions in two accessions, and *BIA1* deletion in one accession, while *CYP702A6* and *AT4G15390* were invariable in copy number.

The two OSCs, *PEN1* and *BARS1*, were located in segments with opposite variation levels. *PEN1* and the neighboring gene *CYP705A1*, both implicated in the arabidiol biosynthesis pathway, were stable in copy number, except for three accessions with full or partial gene deletions: Qui-0 and IP-Deh-1 from Spain and Kyoto from Japan. In the latter, we confirmed partial deletion of both genes by the analysis of its *de novo* genomic assembly (Jiao and Schneeberger, 2020). On the contrary, *BARS1, CYP705A2* and *CYP705A3* were all deleted in several accessions originating from Sweden. We also observed smaller deletions or duplications in this genomic segment, of which the most remarkable was the duplication of *CYP705A2*, detected in 433 (37.6%) accessions. Since the genotypic data for *CYP705A2* and *BARS1* were noisy and indicated more variation than could be revealed by our standard genotyping, we manually inspected short read genomic data which mapped in this region (examples are presented in **Supplemental Figure S10**). In most accessions, *BARS1* lacked the largest intron, where *ATREP11* TE (RC/Helitron super family) is annotated, which might explain lower RD values for *BARS1* compared to other genes (see **Supplemental Figure S3**). Surprisingly, we also observed a mix of reads mapping to *CYP705A2* and *BARS1* loci with and without mismatches, in a large number of accessions. Thus, we called SNPs in the coding sequences of both genes, to get more information on their diversity. Numerous heterozygous SNPs were called in both genes, in the above accessions. Because Arabidopsis is a self-pollinating species and therefore highly homozygous, we hypothesized that the reads with the mismatches originated from duplicated loci, which showed similarity to *CYP705A2* and *BARS1* and mapped to the reference gene models, resulting in heterozygous SNP calls. In support of this hypothesis, we detected heterozygous SNPs at *CYP705A2* locus in 90.6% of accessions with this gene’s duplication, but only in 10.7% of accessions without changes in its copy number (Wilcoxon rank sum test with continuity correction, p.value <2.2×10^−16^; **Supplemental Figure S11A**). Additionally, heterozygous SNPs at *BARS1* locus were present in the same accessions (Pearson’s correlation coefficient r = 0.86; **Supplemental Figure S11B**), although we found only one duplication of *BARS1* with our genotyping methods. We concluded that the sequence differences between *BARS1* and its duplicate prevented its detection by RD or MLPA assays. We also observed low but nonzero read coverage and homozygous SNPs at both loci in some accessions with intermediate RD values for *CYP705A2* (RD_mean_ = 1.5) and *BARS1* (RD_mean_ = 0.6) and with the clear loss of *CYP705A3* (RD_mean_ = 0). In agreement with the gene duplication scenario, this could be explained by the presence of *CYP705A2* and *BARS1* duplicates but absence of the entire region spanning the reference genes *CYP705A2, CYP705A3* and *BARS1*.

To identify the cryptic *BARS1* duplication, we analyzed genomic assemblies of seven accessions: An-1, Cvi-0, Kyoto, Ler-0, C24, Eri-1 and Sha (Jiao and Schneeberger, 2020), four of which have been also genotyped in our study (**Figure 3A**). We re-annotated the entire arabidiol/baruol cluster region in each accession and compared it with the reference (**Supplemental Table S9**). In six accessions, *BARS1* lacked the largest intron, as indicated earlier by short read data (**Supplemental Figure S12**). In Cvi-0, Eri-1 and Ler-0 we identified a nonreference gene, encoding a protein with ∼91% identity to baruol synthase 1 (**Supplemental Figure S13**). In C24, it was also present, but interrupted by ATCOPIA52 retrotransposon insertion, resulting in two shorter ORFs. Based on phylogenetic analysis we concluded that the identified gene was indeed a *BARS1* duplicate and we named it *BARS2* (**Figure 3B**). The differences in the exons of *BARS1* and *BARS2* sequences very well matched the heterozygous SNP positions (**Supplemental Figure S14**). Their introns were much more divergent, which likely affected RD genotyping. Likewise, the probe targeting *BARS1* locus was located in a highly divergent region, which prevented us from detecting this duplication with MLPA.

**Figure 3.**
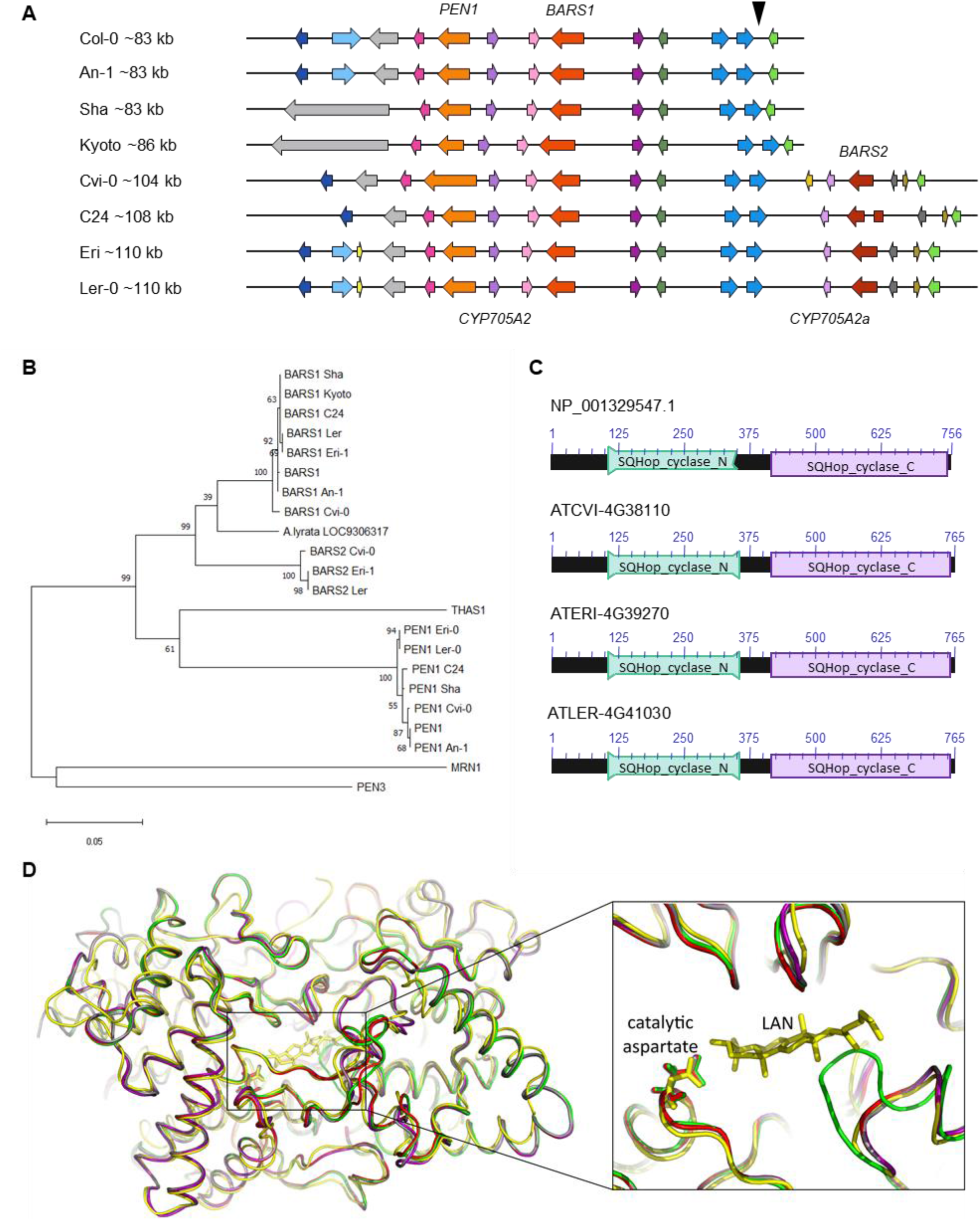
*BARS2* is a *BARS1* duplicate absent from the reference genome and encodes oxidosqualene synthase. A) Organization of arabidiol/baruol gene cluster in Col-0 and seven nonreference accessions. The genomic insertion including *CYP705A2a* and *BARS2* genes is marked with a triangle above the reference cluster. B) Phylogeny of amino acid sequences of clade II OSCs residing in clusters. BARS1 ortholog from A. *lyrata* (LOC9306317) is included. The maximum likelihood tree was generated using the MEGA11 package with Jones-Taylor (JTT) substitution matrix and uniform rates among sites. Values along branches are frequencies obtained from 1000 bootstrap replications. C) Conserved protein domains encoded in *BARS1* (Col-0) and *BARS2* (Cvi-0, Eri-1, Ler-0) genes. SQHop_cyclase_N - squalene-hopene cyclase N-terminal domain (Pfam 13249). SQHop_cyclase_C - squalene-hopene cyclase C-terminal domain (pfam13243) D) 3D models of baruol synthase proteins encoded by *BARS1* and *BARS2*, predicted by ColabFold software, superposed with the crystal structure of human oxidosqualene cyclase in a complex with lanosterol (LAN). The enlargement box highlights the positions of the catalytic aspartate residue in the predicted models. Colors mark superposed models: green (Col-0 BARS1 isoform NP_193272.1), red (Col-0 BARS1 isoform NP_001329547.1), purple (Cvi-0 BARS1 ATCVI-4G38020), grey (Cvi-0 BARS2 ATCVI-4G38110) and yellow (human OSC PDB ID: 1W6K).

The proteins encoded by *BARS2* in Cvi-0, Eri-1 and Ler-0 possessed both N-terminal and C-terminal squalene-hopene cyclase domains, typical for OSCs (**Figure 3C**). We performed three-dimensional (3D) modelling of two reference (Col-0) isoforms of the baruol synthase 1 (the product of *BARS1*) its counterpart from the Relict Cvi-0 as well as a putative baruol synthase 2 (the product of *BARS*2) from Cvi-0, using ColabFol software. Next, we superposed these models with the experimental, crystal structure of human OSC, available in a complex with its reaction product lanosterol (Thoma et al., 2004; Jumper et al., 2021) (**Supplemental information**). All structures were highly similar and we were able to identify potential substrate-binding cavities in the plant enzymes (**Figure 3D; Supplemental Table S10**). Notably, the catalytic aspartate residue D455 present in the human cyclase had its counterparts in the plant OSCs: D493 in the reference isoform NP_193272.1 isoform and D490 in the remaining proteins (**Supplemental file 3**). Together, our data indicated that *BARS2* encoded a novel, so far uncharacterized OSC. As expected, we also found *CYP705A2* duplication in C24, Cvi-0, Eri-1 and Ler-0 assemblies, and we named it *CYP705A2a*. It had 84% identity with *CYP705A2* at the nucleotide level and 88% similarity at the protein level (**Supplemental Figure S15**). *CYP705A2a* and *BARS2* were adjacent to each other and located on the minus strand of the large genomic sequence insertion between *CYP702A6* and *BIA* genes (**Figure 3A**), next to a ∼5 kb long interspersed nuclear element 1 (LINE-1) retrotransposon and some shorter, undefined ORFs. The presence of the insertion increased the size of the entire arabidiol/baruol gene cluster by 21-27 kb.

### Structural diversity of arabidiol/barol gene cluster is associated with the climatic gradient and root growth variation

In the next step, we used the results from the SNP analysis to evaluate the presence/absence variation of both reference (*CYP705A2* + *BARS1*) and nonreference (*CYP705A2a* + *BARS2*) gene pairs in Arabidopsis population (**Supplemental Table S11**). The group with only the reference gene pair present was the largest (PP-AA; 628 accessions). Nearly one-third of the population had both gene pairs (PP-PP; 326 accessions). We also separated two intermediate groups with the local range of occurrence. The first one, with only the nonreference gene pair was found in Azerbaijan, Spain, Bulgaria, Russia, Serbia and U.S.A (AA-PP; 14 accessions). The last group, where we did not detect any of these genes was mostly observed at Bothnian Bay coast collection site in North Sweden (AA-AA; 15 accessions). For 73 accessions, the data were unconclusive. The accuracy of group assignments was validated by sequence analysis of *de novo* genomic assemblies for An-1, Kyoto, Mitterberg-2-185 and Kn-0 (PP-AA group) as well as Cvi-0, Ler-0, Dolna-1-40 and Ty-1 (PP-PP group). Additionally, the results of PCR amplification with gene-specific primers and genomic DNA template for a subset of 36 accessions from all four groups confirmed the differences between them (**Supplemental Figure S16**). We could not detect *BARS2*-specific product in many samples from the AA-PP group, however we did detect the band for *CYP705A2a*. We suppose that the *BARS2* sequence might further diverge in this minor group.

The accessions with the nonreference gene pair (AA-PP; PP-PP) dominated among Relicts (81%), Spain (60%) and Italy/Balkan/Caucasus (89.6%) genetic groups, but constituted the minority at the north and east margins of the species range (North Sweden 18.6%, South Sweden 16%, Asia 9.4%; **Figure 4B**). They were also mostly absent among U.S.A accessions. The widespread presence of *CYP705A2a* and *BARS2* genes in Relicts suggested that the duplication event preceded the recent massive species migration, which took place in the post-glacial period and shaped the current Arabidopsis population structure (Lee et al., 2017). We next visualized the four groups in the principal component analysis (PCA) plots, generated with genome-wide biallelic SNPs (1001 Genomes Consortium, 2016; Zmienko et al., 2020). At low linkage disequilibrium parameter, where the contribution of the ancestral alleles to PCA was highest, there was a clear convergence of the PC1 and PC2 components with the presence /absence of the gene duplication (**Figure 4C; Supplemental Figure S17**). This suggested that the presence/absence of the genomic insertion containing *CYP705A2a* and *BARS2* genes had some impact on the current geographic distribution of Arabidopsis accessions. We then evaluated the accessions’ latitudes of origin and found that accessions with the nonreference gene pair originated from significantly lower latitudes, compared to the remaining ones (one-way rank-based analysis of variance ANOVA, p.value<0.001, followed by Dunn’s test with BH correction, p.value<0.001) (**Figure 4D**). This difference was noticeable even within individual countries and it was significant for Germany, Spain and Italy (**Supplemental Figure S18**). We observed the reverse trend in Russia, where PP-AA accessions were in great excess (88%) and in France, however, we also noticed that PP-AA accessions outnumbered PP-PP accessions in the Pyrenees, Alps and Tian Shan mountain ranges (**Supplemental information**). It suggested there was association between arabidiol/baruol gene cluster variation and environmental conditions, therefore, we decided to check it in the next step. Since the climate is a substantial selection factor, we also checked for the phenotypic variability between the most abundant PP-AA and PP-PP groups. To this end, we performed two-group comparisons of 516 phenotypic and climatic variables retrieved from the Arapheno database (Seren et al., 2017; Togninalli et al., 2020) and focused on these which significantly differed between both groups (Wilcoxon rank sum test with continuity correction, p.value <0.05) (**Supplemental Table S12**). Prominently, we observed differences for 88 climatic variables (Exposito-Alonso et al., 2019), especially maximal and minimal temperature conditions, precipitation and evapotranspiration (**Figure 5A**). Apart from the climate data, 40 diverse phenotypes varied significantly between both groups. Although some of these differences, e.g. flowering-related phenotypes, might be influenced by another genetic factor, independent from the arabidiol/baruol gene cluster structure (Li et al., 2010), we paid special attention to root growth-related phenotypes, since all Arabidopsis MGCs are considered to have root-specific expression (Huang et al., 2019). We observed significant differences between PP-AA and PP-PP groups in root growth dynamics, which was analyzed during the first week after germination by Bouain et al. (2018). More specifically, roots of PP-PP accessions elongated slower compared to PP-AA accessions (**Figure 5B**). Additionally, PP-PP accessions showed a significantly lower rate of root organogenesis from explants under one of three growth conditions tested in another study (Lardon et al., 2020) (**Figure 5C**). We next applied a linear mixed model in a Genome-Wide Association Study on the same 516 phenotypes, in order to independently evaluate the significance of our observations, after correction for the population structure and multiple testing. We used a genome-wide matrix of over 250 thousand biallelic SNPs, supplemented with a SNP-like encoded information about the gene duplication status (only PP-AA and PP-PP groups were analyzed). Although the association of *CYP705A2a* and *BARS2* presence/absence variation was not statistically significant for any variable we tested, we again obtained lowest p-values for the climatic data and root organogenesis phenotypes (**Figure 5D, Supplemental Table S12**). We then checked for the genetic interactions between thalianol and arabidiol/baruol clusters, to exclude the possibility that they affected our results, since the distribution of discontiguous and compact versions of thalianol gene cluster was also strongly associated with the latitude (**Supplemental Figure S19**). However, the variation in thalianol gene cluster organization did not explain the variation in root growth phenotypes, contrary to structural variation of the arabidiol/baruol gene cluster.

**Figure 4.**
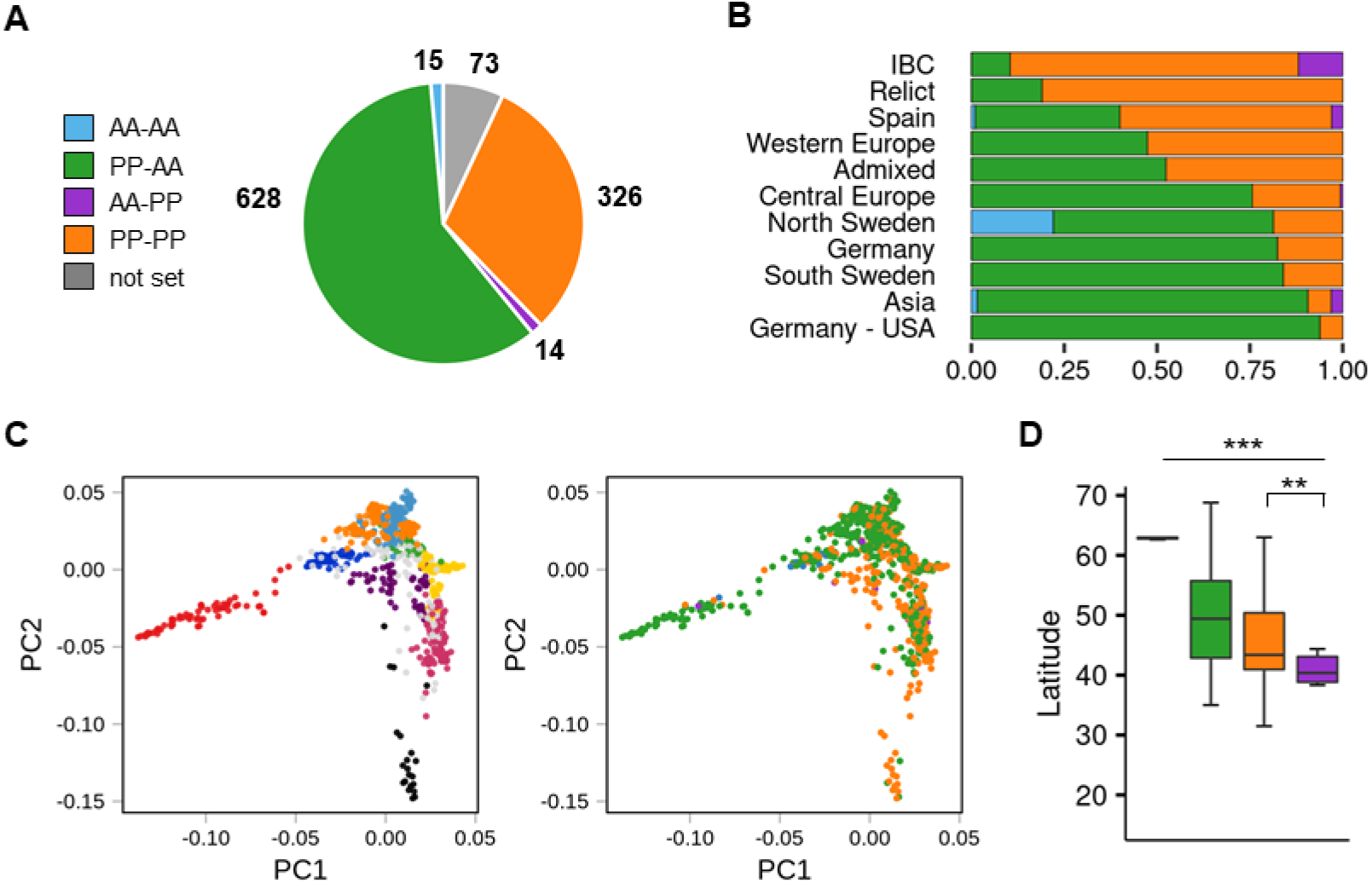
Population-scale diversity of *CYP705A2* and *BARS1* duplication status. A) The sizes of four groups differing by the presence (PP)/absence (AA) of *CYP705A2*-*BARS1* and *CYP705A2*-*BARS2* gene pairs. B) Group distribution among the genetic groups. U.S. accessions from the German group were separated from the remaining accessions. C) Principal component analysis (PCA) plots, generated at linkage disequilibrium LD = 0.3. The first two components are presented. Accessions are colored according to their genetic group (left) or CYP-BARS status (right). U.S. accessions were not included in the analysis, in order to better visualize the remaining groups. PCA plots with other LD parameters are **in Supplemental Figure S17** D) Latitudes of accessions’ sites of origin, grouped by CYP-BARS status. One-way rank-based analysis of variance ANOVA, p.value<0.001, followed by Dunn’s test with BH correction, **p.value<0.05 (PP-PP vs AA-PP); ***p.value < 0.001 (all the other pairwise comparisons).

**Figure 5.**
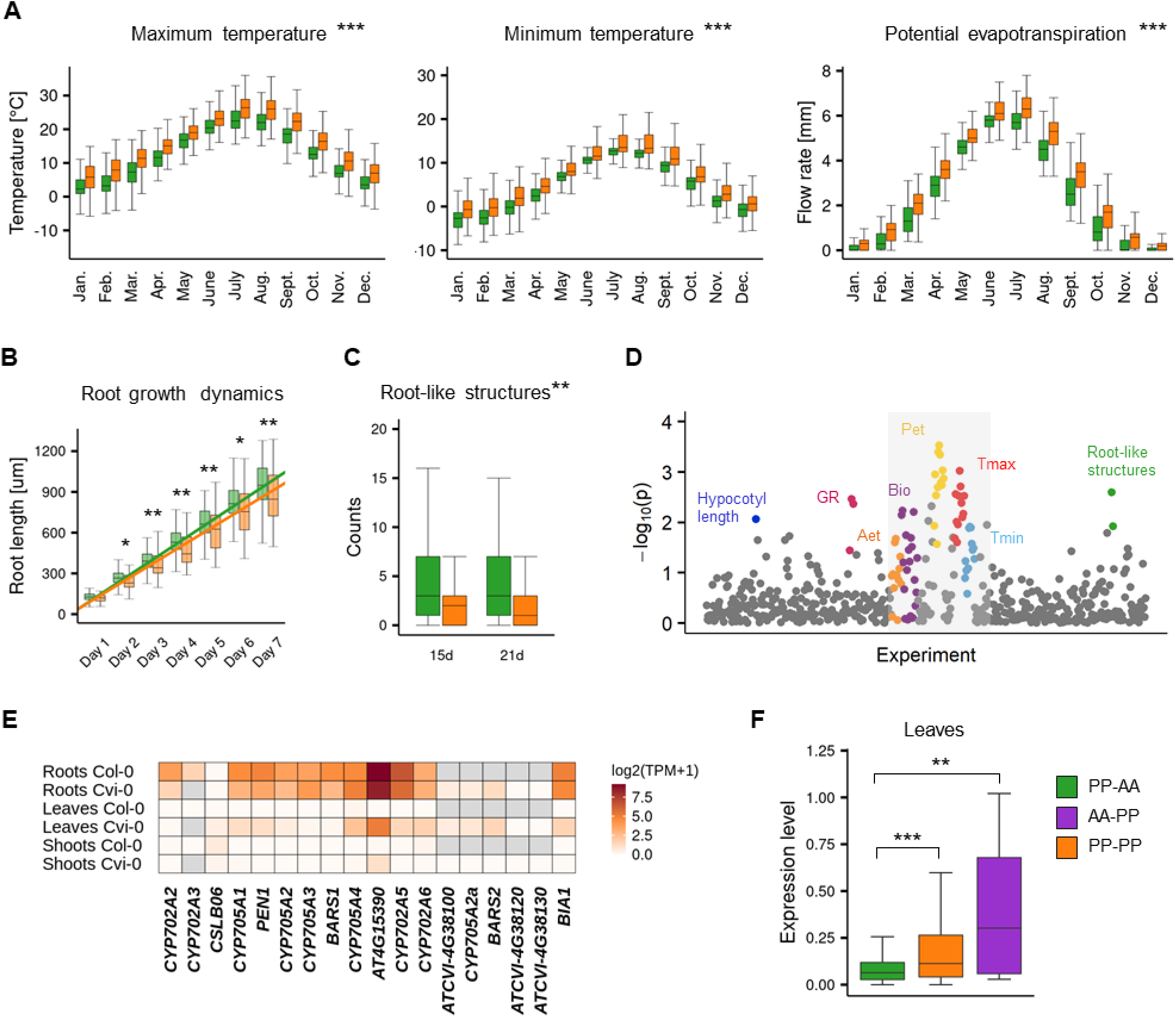
Phenotypic variation of PP-AA and PP-PP groups. A-C) Two-group comparisons of climatic (A), root growth dynamics (B) and root organogenesis (C) data between PP-AA (green) and PP-PP (orange) accessions. Stars denote the significance of Wilcoxon rank sum test with continuity correction, *p.value<0.1, **p.value<0.05, ***p.value<0.001. D) Results of a genome-wide association study for PP-AA / PP-PP allelic variation. Study with climatic data is in the grey box E) Tissue specificity of arabidiol/baruol gene cluster expression in Col-0 and Cvi-0. F) Population-level differences in gene expression in leaves among the PP-AA, PP-PP and AA-PP groups. Expression levels are shown as log_2_(TPM+1). Stars denote the significance of one-way rank-based analysis of variance ANOVA, p.value<0.001, followed by Dunn’s test with BH correction, **p.value<0.05, ***p.value<0.001. Source data are available in the Arapheno database (plots A-C), Supplemental Table S12 (plot D) and Supplemental Table S13 (plots E-F).

In the reference accession Col-0, all genes in the arabidiol/baruol cluster are low-expressed and active almost exclusively in roots (**Supplemental Figure S20A**). In search of the possible links between arabidiol/baruol gene cluster structure and phenotypic variation, we investigated *CYP705A2, BARS1, CYP705A2a* and *BARS2* expression profiles in Col-0 and Cvi-0. We used RNA-Seq data from roots, shoots and leaves, which we retrieved from the studies where these accessions were grown in parallel under standard conditions (Kawakatsu et al., 2016; van Veen et al., 2016). We mapped the data to the respective (Col-0 or Cvi-0) annotated genome and compared the gene expression profiles (**Figure 5E; Supplemental Table S13**). In both accessions, the arabidiol/baruol gene cluster was silenced in shoots, except for the low activity of acyltransferase gene *AT4G15390*, detected in Cvi-0. Also in both accessions, the clusters were active in roots, and the expression of *AT4G15390* was much stronger than the remaining genes. In Cvi-0, genes located in the genomic insertion (*CYP705A2a, BARS2* and *ATCVI-4G38100* which encodes a protein with partial similarity to acyltransferase) were also expressed, although at a lower level, compared to the rest of the cluster. Surprisingly, in leaves of Cvi-0, but not Col-0, we also detected transcriptional activity within arabidiol/baruol gene cluster. Most clustered genes were expressed at lower level than in Cvi-0 roots, and the transcripts of *CYP705A2, CYP705A3* and *BARS1* were barely detectable. However, *ATCVI-4G38100, CYP705A2a* and *BARS2* had similar expression in leaves and roots. Taking these observations into account, it should not be excluded that the metabolic products of arabidiol/baruol gene cluster activity in the roots and leaves of Cvi-0 accession are not identical.

Since PP-PP group represented a substantial fraction of the Arabidopsis population, we wanted to check whether the gene expression profile, which we observed in leaves of Cvi-0 was ubiquitous among the accessions from this group. To this end, we analyzed RNA-Seq data for 552 accessions, mapped against the reference genome (Kawakatsu et al., 2016) and we compared the *BARS1* expression level between AA-PP, PP-PP and PP-AA groups. It was significantly higher in accessions with *CYP705A2a* + *BARS2* gene pair, compared to the PP-AA group (one-way rank-based analysis of variance ANOVA, p.value<0.001, followed by Dunn’s test with BH correction, p.value <0.05) (**Figure 5F**), in agreement with our predictions that *BARS2* was expressed in the leaves of these accessions and that reads from *BARS2* transcripts mapped to *BARS1* locus, elevating its measured expression level. We also re-mapped the raw RNA-Seq reads from Ty-1 and Cdm-1 (PP-PP group), as well as Kn-0 and Sha (PP-AA group) accessions to their respective genomic assemblies and separately measured the expression levels of *BARS1* and *BARS2*. As expected, in the leaves of PP-PP accessions, *BARS2* was expressed, while *BARS1* was not (**Supplemental Figure S20B**).

### Paired terpenoid synthase and cytochrome P450 genes are more variable than non-paired ones

In many plant genomes, genes encoding terpenoid synthases (TSs, this includes OSCs analyzed in our study) are positioned in the vicinity of CYPs more often than expected by chance. Therefore, they frequently exist as TS-CYP pairs (Boutanaev et al., 2015). TS-CYP pairs located in MGCs had similar (either high or low) copy number diversity and were frequently duplicated or deleted together. We wanted to check whether this observation could be extended to other TS-CYP pairs in the Arabidopsis genome. Therefore, we created a comprehensive list of 48 TSs and 242 CYPs, based on trusted sources (Paquette et al., 2000; Bak et al., 2011; Nelson and Werck-Reichhart, 2011; Boutanaev et al., 2015). We then retrieved information about each gene’s copy number diversity among 1,056 accessions (**Supplemental Tables S14-S15**). For 13 TSs, including *THAS1* and *BARS1*, we observed gains or losses in at least 1% of accessions. Only 33 CYPs showed such variability and they represented three clans: CYP71 (26 variable genes out of 151), CYP85 (6 variable genes out of 29) and CYP72 (1 variable gene out of 19). The remaining clans showed very low variability. Next, for each TS, we selected all CYPs within +/- 30-kb distance, which produced 38 pairs between 18 TSs and 27 CYPs, including pairs in thalianol, marneral, tirucalladienol and arabidiol/baruol gene clusters, as well as other putative secondary metabolism clusters, listed in plantiSMASH resource (Kautsar et al., 2017). Subsequent group comparisons revealed that TSs and CYPs occurring in pairs were more variable than their non-paired counterparts (Wilcoxon rank sum test with continuity correction, p.value<0.01 for TSs, p.value<0.001 for CYPs). Moreover, for numerous TS-CYP gene pairs, the CNV rates and patterns of both genes were similar (**Supplemental Figure S21**). This indicated than nonrandom association of TSs and CYPs is preserved in Arabidopsis genome by physical constraints or selection.

## Discussion

According to our current understanding of the MGC formation phenomenon, nonrandom gene clustering in eukaryotes is linked with the highly dynamic chromosomal regions. Numerous studies highlighted that structural variations are the main genetic drivers of metabolic profiles diversity and MGC evolution in plants (Fan et al., 2020; Li et al., 2020; Liu et al., 2020a; Liu et al., 2020b; Zhan et al., 2020; Katz et al., 2021). These studies suggested that plant MGCs are dynamically evolving and the genetic mechanisms that originally lead to their formation may be captured at the intraspecies genetic variation level. Similar conclusions were drawn from the previous study of a filamentous fungus *Aspergillus fumigatus*, in which the secondary metabolic pathway genes are commonly organized into clusters (Lind et al., 2017). During the evolution, new biochemical pathways are tuned and tested by many rounds of natural selection. The analysis of the intraspecies MGC variants, which are more recent than the variants found in interspecies comparisons, may provide an important insight into the formation of the clustered gene architectures and the plant metabolic diversity, in a small evolutionary time frame.

The four MGCs in Arabidopsis are implicated in the biosynthesis of structurally diverse triterpenes and are dated after the α whole-genome duplication event, which occurred in Brassicaceae lineage ∼23-43 Mya (Field et al., 2011). These MGCs are assembled around the gene(s) encoding clade II OSCs. It has been shown that in various *Brassicaceae* genomes, clade II OSCs are often colocalized with the genes from CYP705, CYP708 and CYP702 clans, and with the genes from the acyltransferase IIIa subfamily (Liu et al., 2020b). Bioinformatic studies also revealed that TSs and CYPs are paired in plant genomes more frequently than expected (Boutanaaev et al., 2015). We found that in Arabidopsis, the physical proximity of CYPs and TSs was associated with increased CNV rates for these genes, compared to the non-paired ones. It might suggest that the occurrence of such a specific gene mix, combined with the structural instability of its genomic neighborhood, boosted the potential to produce novel metabolic pathways. The four Arabidopsis MGCs had different levels of variation, which generally reflected the phylogeny of clade II OSCs contained in these clusters. Of them, MRN1 is most divergent in amino acid sequence. It is also mono-functional, i.e., catalyzes the formation of one specific product - marneral (Xiong et al., 2006). Functional studies indicated a critical role of marneral synthase in Arabidopsis development (Go et al., 2012). Consistent with these findings, *MRN1* was the only clustered OSC gene, which was not affected by deletions or duplications, in any accession. Also, the neighboring CYPs were stable in copy number. Our results indicate that marneral gene cluster is fixed in Arabidopsis genome.

The arabidiol/baruol gene cluster was the most variable MGC. It comprises few gene subfamilies, but is significantly expanded compared to the sister species *A. lyrata*, which is suggestive of recent duplications. For example, *PEN1* and *BARS1* have only one ortholog in *A. lyrata, LOC9306317*. Accordingly, we observed an exceptionally high rate of intraspecific gene gains and losses within this MGC. The segmentation of arabidiol/baruol gene cluster into variable and invariable gene blocks may result from the ongoing process of selection-driven fixation of the arabidiol sub-cluster. The products of *PEN1* and *CYP705A1* are involved in response to jasmonic acid treatment and infection with the root-rot pathogen *Pythium irregulare* (Sohrabi et al., 2015). Moreover, arabidiol may be further converted to arabidin, in the pathway involving acyltransferase encoded by *AT5G47950*, which is located in thalianol gene cluster (Huang et al., 2019) and was also invariable in copy number. The fixation of genes involved in the arabidin biosynthesis may indicate the biological significance of this pathway. Crispr mutants with disrupted *AT5G47950* gene had significantly shorter roots than the wild-type plants and arabidin did not accumulate in these roots (Bai et al., 2021). Interestingly, *A. lyrata* is able to convert apo-arabidiol (the product of arabidiol degradation) into downstream compounds, despite the lack of arabidiol synthase (Sohrabi et al., 2017). This indicates there may be modularity of the biosynthetic pathways in plants. It might facilitate assembling a biosynthesis network and lead to increase in the repertoire of the secondary metabolites produced by the plant. Understanding the complexity of this network may be supported by in-depth analysis of MGC intraspecies variation.

The initial diversity of 2,3-oxidosqualene cyclization products generated by the plant is determined by OSCs diversity. Here we report the discovery of *BARS2* gene, which was found in numerous accessions, but was absent from Col-0, hence it was absent from the reference genome. Our data indicated that *BARS2* encodes a functional clade II OSC. It is worth to notice that baruol synthase 1 encoded by its closest paralog, *BARS1*, has the lowest product specificity among plant OSCs (Lodeiro et al., 2007; Ghosh et al., 2016). Why some OSCs are highly multi-functional is not well understood. It has been suggested that they are undergoing evolution towards increased product specificty. It was demonstrated that only two amino acid changes in the cycloartenol synthase lead to its conversion into an accurate lanosterol synthase (Lodeiro et al., 2005). Biochemical characterization of baruol synthase 2 and its comparison with baruol synthase 1 may help reveal the role of particular amino acids in acquiring specificity for given products.

According to our data, *BARS2* and *CYP705A2a* gene pair may be present in nearly one-third of Arabidopsis population and their presence/absence variation is associated with the climatic gradient and the root growth dynamics. Additional studies are needed to assess whether these observations may be linked to the expression of these two genes or to the differences in transcriptional activity of the entire cluster (Wegel et al., 2009; Yu et al., 2016; Roulé T). In Col-0, MGCs are embedded in local hotspots of three-dimensional chromatin interactions. Their activation in roots and repression in leaves is combined with the distinct chromatin condensation states and nuclear repositioning of MGC regions between these tissues (Nützmann et al., 2020). Loss of the histone mark H3K27me3 in *clf/swn* mutant resulted in the loss of interactive domains associated with the thalianol, marneral and arabidiol/baruol cluster regions, indicating that different transcriptional states of these MGCs are strictly regulated by the switches in their conformation. Curiously, in accessions with *CYP705A2a* and *BARS2*, we observed some transcriptional activity of arabidiol/baruol cluster genes in leaves. The presence of ∼25-kb insertion in the arabidiol/baruol gene cluster may alter its structure and affect the epigenetic regulation of its activity.

Thalianol gene cluster was the second most variable MGC in our analysis. The first evidence for its structural diversity comes from the study of Liu et al. (2020a), who found large deletions affecting thalianol biosynthesis genes in ∼2% of the studied accessions. Since our approach was specifically focused on CNV analysis and was duplication-aware, we were able to detect over two times more CNVs in a similar population (4.7%), with 49 accessions carrying gene deletions and five accessions with gene duplications. Apart from the identification of two new variants – one large deletion and a duplication, we validated earlier assumptions that the nonreference compact version of the thalianol gene cluster is predominant in Arabidopsis. Moreover, it is also better conserved, compared to the discontiguous version. It remains to be investigated whether the tighter clustering of thalianol gene cluster may be advantageous in certain environmental conditions or it is just less prone to structural variation, due to physical constraints.

Triterpenes are high-molecular-weight non-volatile compounds, which are likely to act locally. However, they may be further processed and generate various breakdown products, both volatile and non-volatile, which may be biologically active (Sohrabi et al., 2015; Sohrabi et al., 2017). The compounds of plant origin may be also metabolized by plant-associated microbiota. A recent study demonstrated that various coctails of thalianin, thalianyl fatty acid esters and arabidin attracted or repelled various microbial communities present in the soil, and participated in the plant’s active selection of root microbiota (Huang et al., 2019). In fact, a small but significant effect of Arabidopsis genotype on the root microbiome has been demonstrated previously (Bulgarelli et al., 2012; Lundberg et al., 2012). In a recent study by Karasov et al. (2022), bacterial communities that colonized the leaves of 267 local Arabidopsis populations, assessed at various localizations in Europe, formed two distinct groups, strongly associated with the latititude. Specifically, a significant latitudinal cline was observed for the strains of *Sphingomonas* genus, which is commonly associated with Arabidopsis (Bodenhausen et al., 2013). Various *Sphingomonas* species possess a range of biodegradative and biosynthetic capabilities (Mohn et aal., 1999; Asaf et al., 2020). *Sphingomonas* is implicated in promoting Arabidopsis growth and increasing drought resistance, and protecting the plants against the leaf-pathogenic *Pseudomonas syringae* (Innerebner et al., 2011; Luo et al., 2019). Prominently, in the study by Karasov et al. (2022), the host plant genotype alone could explain 52% to 68% of the observed variance in the phyllosphere microbiota. Moreover, the microbiome type was strongly associated with the index of the dryness of the local environment, based on recent precipitation and temperature data. We propose that the genetic diversity of terpenoid metabolism pathways in Arabidopsis may be interdependent on the diversity of soil bacterial communities present in various environments, and this relationship might play role in Arabidopsis adaptation to climate-driven selective pressures. Further exploration of MGC diversity may help us understand these biotic interactions.

Currently, the bioinformatic identification of new MGC candidates is mainly based on the combination of physical gene grouping and their coexpression. The accuracy and sensitivity of such approaches strongly depend on the abundance of data from various tissues, time points, and environmental conditions (Wisecaver et al., 2017). We suggest that the analysis of intraspecies genetic and transcriptomic variation may provide a valuable addition to MGC studies. The genome of one individual may not be representative enough to reveal the entire complexity of a given pathway, not to mention the metabolic diversity of the entire species (Kawakatsu et al., 2016; Shirai et al., 2017; Zmienko et al., 2020; Katz et al., 2021). With the rapid increase in the number of near-to-complete assemblies of individuals’ genomes facilitated by the development of third-generation sequencing technologies, we are now entering the era of intense exploration of the impressive plasticity of plant metabolic pathways.

## Methods

### Plant material and DNA samples

Arabidopsis seeds were obtained from The Nottingham Arabidopsis Stock Centre. The seeds were surface-sterilized, vernalized for 3 days, and grown on Jiffy pellets in ARASYSTEM containers (BETATECH) in a growth chamber (Percival Scientific). A light intensity of 175 mmol m^-2^ s^-1^ with proportional blue, red, and the far red light was provided by a combination of fluorescent lamps (Philips) and GroLEDs red/far red LED Strips (CLF PlantClimatics). Plants were grown for 3 weeks under a 16-h light (22°C)/8-h dark (18°C) cycle, at 70% RH, nourished with half-strength Murashige & Skoog medium (Serva). Genomic DNA for MLPA and ddPCR assays was extracted from 100 mg leaves with a DNeasy Plant Mini Kit (Qiagen), according to manufacturer’s protocol, which included RNase A treatment step.

### RD assays

To determine the boundaries of each MGC, the relevant literature and gene coexpression datasets were surveyed (Fazio et al., 2004; Xiong et al., 2006; Xiang et al., 2006; Lodeiro et al., 2007; Field and Osbourn, 2008; Morlacchi et al., 2009; Field et al., 2011; Go et al., 2012; Thimmappa et al., 2014; Sohrabi et al., 2015; Yasumoto et al., 2016; Wisecaver et al., 2017). TAIR10 genome version and Araport 11 annotations (Cheng et al., 2017) were used as a reference in all analyses. Short read sequencing data from Arabidopsis 1001 Genomes Project (1001 Genomes Consortium, 2016) were downloaded from National Center for Biotechnology Information Sequence Read Archive repository (PRJNA273563), processed and mapped to the reference genome as described in (Zmienko et al., 2020). The gene copy number estimates based on read-depth analysis of short reads (RD dataset) were generated previously and are available at http://athcnv.ibch.poznan.pl. Accessions BRR57 (ID 504), KBS-Mac-68 (ID 1739), KBS-Mac-74 (ID 1741) and Ull2-5 (ID 6974), which we previously identified as harboring unusually high level of duplications, were removed from the analysis.

### MLPA assays

MLPA probes were designed according to a procedure designed previously and presented in detail in (Samelak-Czajka et al., 2017). Probe genomic target coordinates are listed in **Supplemental Table S16**. The MLPA assays were performed using 5 ng of DNA template with the SALSA MLPA reagent kit FAM (MRC-Holland). The MLPA products were separated by capillary electrophoresis in an ABI Prism 3130XL analyzer at the Molecular Biology Techniques Facility in the Department of Biology at Adam Mickiewicz University, Poznan, Poland. Raw electropherograms were quality-checked and quantified with GeneMarker v.2.4.2 (SoftGenetics), with peak intensity and internal control probe normalization options enabled. Data were further processed in Excel (Microsoft). To allow easy comparison of RD and MLPA values, the MLPA results were normalized to a median of all samples’ intensities and then multiplied by 2, separately for each gene/MLPA probe.

### ddPCR assays

Genomic DNA samples were digested with XbaI (Promega). DNA template (2.5 ng) was mixed with 1× EvaGreen ddPCR Supermix (Bio-Rad), 200 nM gene-specific primers (**Supplemental Table S17**) and 70 μl of Droplet Generation Oil (Bio-Rad), then partitioned into approximately 18,000 droplets in a QX200 Droplet Generator (Bio-Rad), and amplified in a C1000 Touch Thermal Cycler (Bio-Rad), with the following cycling conditions: 1× (95 °C for 5 min), 40× (95 °C for 30 s, 57 °C for 30 s, 72 °C for 45 s), 1× (4 °C for 5 min, 90 °C for 5 min), with 2 °C/s ramp rate. Immediately following end-point amplification, the fluorescence intensity of the individual droplets was measured using the QX200 Droplet Reader (Bio-Rad). Positive and negative droplet populations were automatically detected by QuantaSoft droplet reader software (Bio-Rad). For each accession and each gene, the template CNs [copies/μl PCR] were calculated using Poisson statistics, background-corrected based on the no-template control sample and normalized against the data for previously verified non-variable control gene *DCL1*.

### PCR assays

Genomic DNA samples (5 ng) were used as templates in 20 μl reactions performed with PrimeSTAR GXL DNA Polymerase (TaKaRa), according to the manufacturer’s instructions, in a three-step PCR. Amplicons (10 ul) were analyzed on 1% agarose with 1kb Gene Ruler DNA ladder (Fermentas). Primer sequences are listed in **Supplemental Table S17**. Primer pairs for *BARS1-BARS2* and *CYP705A2-CYP705A2a* were designed in corresponding genomic regions, that assured primer divergence between the paralogs. However, primers designed for *CYP705A2* produced unspecific bands of ∼5kb in many samples. Therefore, this gene was excluded from the analysis.

### Genotype assignments

For MLPA dataset, genotypes were assigned to each gene and each accession based on normalized MLPA values of ≤1 for LOSS genotype and >3 for GAIN genotype. The remaining cases were assigned REF genotype. For RD dataset, the respective RD thresholds were ≤1 for LOSS genotype and >3.4 for GAIN genotype, except for BARS1, for which both thresholds were lowered by 0.2. The remaining cases were assigned REF genotype. For ddPCR, genes with normalized CN=0 were assigned LOSS genotype and genes with normalized CN=2 were assigned REF genotype. The RD, MLPA and ddPCR datasets were then combined using the following procedure. For genes and accessions covered by multiple datasets, the final genotype was assigned based on all data.

Discordant genotype assignments (21 out of 1,784 covered by multiple datasets) were manually investigated and 19 of them were resolved (**Supplemental Figure S4; Supplemental Table S7**). Out of the remaining 32,000, which were assayed with one method only, the genotype was manually corrected in 13 cases with values very close to the arbitrary threshold, based on population data distribution. Final genotype assignments for each gene and each accession are listed in **Supplemental Table S6**.

### Sanger sequencing

The genomic DNA of Mir-0 accession (ID 8337) was used as a template (2 ng) for amplification using PrimeSTAR® GXL DNA Polymerase (TaKaRa), in a 40-µl PCR reaction with 0.3 µM primers OP009 and OP010, according to general manufacturer instructions. The amplified product, of ∼8 kb in length, was purified with DNA Clean & Concentrator (ZYMO Research) and checked by gel electrophoresis and analysis on NanoDrop™ 2000 Spectrophotometer. The purified product (110 ng) was mixed with 1 ul of sequencing primer Mar02_R and sequenced on ABI Prism 3130XL analyzer at the Molecular Biology Techniques Facility in the Department of Biology at Adam Mickiewicz University, Poznan, Poland. Sequencing files were analyzed with Chromas Lite v. 2.6.6. (Technelysium) software.

### De novo genomic assemblies generation, annotation and analysis

Mitterberg-2-185 and Dolna-1-40 genomic sequences were extracted, sequenced on 1 MinION flowcell (*Oxford Nanopore Technologies*) each and assembled *de novo* with Canu. Genomic sequences of interest (corresponding to thalianol gene cluster for Mitterberg-2-185 and tirucalladienol gene cluster for Dolna-1-40) were then retrieved with megablast (blast-2.10.0+ package) using TAIR10 reference genomic sequence as a query. The remaining de novo assemblies were retrieved from the following public databases. The PacBio-based genomic assemblies, gene annotations and orthogroups for An-1, C24, Cvi-0, Eri-1, Kyoto, Ler-0 and Sha accessions, as well as the reference genome coordinates of the hotspots of rearrangements, were downloaded from Arabidopsis 1001 Genomes Project Data Center (MPIPZJiao2020) or retrieved from the corresponding paper (Jiao and Schneeberger, 2020). Assembled genomic sequences of Ty-1 (PRJEB37258), Cdm-0 (PRJEB40125) and Kn-0 (PRJEB37260) accessions were retrieved from NCBI/Assembly database (Sayers et al., 2022). Gene prediction was performed with Augustus v.3.3.3 (Stanke and Morgenstern, 2005) with the following settings: “Species *Arabidopsis thaliana”*, “both strands”, “few alternative transcripts” or “none alternative transcripts”, “predict only complete genes”. These parameters were first optimized by gene prediction in the corresponding TAIR 10 genomic sequence and comparison with Araport 11 annotation. For previously annotated assemblies, we added information about the newly predicted genes to existing annotations. The protein sequences of *de novo* predicted genes and the information about their best blast hit in the reference genome are available in **Supplemental information**. The search for conserved domain organization was performed with the online NCBI search tool against Pfam v.33.1 databases. Protein sequence alignment was done with Multalin or EMBL online tools (Corpet, 1988; Madeira et al., 2019). TEs were annotated with RepeatMasker software version 4.1.2 (http://www.repeatmasker.org), using homology-based method with TAIR10-transposable-elements reference library.

### Identification of chromosomal inversions

The BreakDancerMax program from the BreakDancer package v.1.3.6 (Chen et al., 2009) was used to detect inversions in each of 997 samples with paired-end data and unimodal insert size distribution. Variants were called separately for each accession and each chromosome. Only calls with lengths within the range 0.5 kbp – 50 kbp and with the Confidence Score >35 were retained.

Since BreakDancerMax output included numerous overlapping calls for individual accessions, we first minimized its redundancy. From the overlapping regions, we kept one variant with i) the highest Confidence Score, and ii) the highest number of supporting reads. If two or more overlapping variants had the same score and the number of supporting reads number, maximized coordinates of these variants were used. This step was carried out in two iterations, considering the 50% reciprocal overlap of the variants. Then, the inversions that overlapped with the thalianol gene cluster were selected from each genome-wide dataset.

### SNP calling at *CYP705A2* and *BARS1* genes

Variants (SNPs and short indels) were called with DeepVariant v.1.3.0 in WGS mode and merged with GLnexus (Yun et al., 2021). Analysis was performed for *CYP705A2* and *BARS1* genomic loci. The results were further filtered to include only biallelic variants, that were located in the exons of each gene (for *BARS1*, exon intersections from two transcript models were used). The number of heterozygous positions was then calculated for each accession and each gene. The same procedure was repeated by taking into account only biallelic variants with at least 1% frequency, which resulted in nearly identical results. Both types of analysis led to the selection of the same set of accessions with duplication at both loci.

### Genome-wide SNP analysis

Variants for 983 accessions with known *CYP705A2* + *BARS1 & CYP705A2a* + *BARS2* pair status were downloaded from the 1001 Genomes Project Data Center (1001genomes_snp-short-indel_only_ACGTN_v3.1.vcomparedwithsnpeff file) (1001 Genomes Consortium, 2016). Data preprocessing was performed using PLINK v.1.90b3w (https://www.cog-genomics.org/plink/1.9/; Chang et al., 2015). Variants with missing call rates exceeding value 0.5 and variants with minor allele frequency below 3% were filtered out. The LD parameter for linkage disequilibrium-based filtration was set as follows: indep-pairwise 200’kb’ 25 0.5. Over 250,000 SNPs were used for PCA analysis with EIGENSOFT v.7.2.1 (Price et al., 2006; Patterson et al., 2006). PCA for a wide LD range between 0.3 - 0.9 was calculated. U.S.A accessions which only recently separated geographically from the rest of the population (Lee et al., 2017) were excluded, to ensure better visibility of the remaining accessions. The ggplot2 package was used for data visualization in R v4.0.4 (https://www.r-project.org; Wickham, 2016).

### Genome-Wide Association Study and phenotype analysis

The entire set of 516 phenotypes from 26 studies was downloaded from the Arapheno database on 26 April 2022 (Seren et al., 2017; Togninalli et al., 2020). The above genome-wide SNP dataset, to which we added a biallelic variant representing PP-AA or PP-PP group assignment, was used. The IBS kinship matrix was calculated on 954 accessions. Association analysis was performed for each phenotype using a mixed model correcting for population structure using Efficient Mixed-Model Association eXpedited, version emmax-beta-07Mar2010 (Kang et al., 2008). Input file generation and analysis of the results were performed with PLINK v.1.90b3w and R v4.0.4.

### Analysis of RNA-Seq data

Processed RNA-seq data from leaves for 728 accessions (552 in common with our study) mapped to the reference transcriptome (Kawakatsu et al., 2016) were downloaded from NCBI/SRA (PRJNA319904), normalized and used to compare *BARS1* expression levels between PP-AA, PP-PP and AA-PP groups. Additionally, raw RNA-Seq reads from leaves were downloaded from the same source for accessions-specific mapping and analysis of Cdm-0, Col-0, Cvi-0, Kn-0, Ty-1 and Sha accessions. Raw RNA-Seq reads from roots and shoots of Col-0 and Cvi-0 accessions were retrieved from BioProject PRJEB14092 (van Veen et al., 2016). SRA Toolkit v2.8.2. (https://github.com/ncbi/sra-tools) and FastQC v0.11.4 (https://www.bioinformatics.babraham.ac.uk/projects/fastqc/) were used for downloading the raw reads and for the quality analysis. For Cdm-0, Kn-0 and Ty-1 genomes, .gtf files were generated based on Augustus results, that included the annotations for the genes of interest. (provided as **Supplemental file 4**). Raw reads were mapped to the respective genomes using the STAR aligner version 2.7.8a (Dobin et al., 2013). STAR indices were generated with parameters: “--runThreadN 24 --sjdbOverhang 99 --genomeSAindexNbases 12”. The following parameters were used for the mapping step: “--runThreadN 24 --quantMode GeneCounts --outFilterMultimapNmax 1 --outSAMtype BAM SortedByCoordinate --outSAMunmapped Within”. Bioinfokit v1.0.8 https://zenodo.org/record/3964972#.Yyw6oRzP1hE) was used to convert .gff3 to .gtf files. Transcripts per million (TPM) values and fragments per kilobase exon per million reads (FPKM) with total exon length for each gene were computed in R v4.0.4.

### Analysis of TS-CYP pairs

A list of Arabidopsis CYP genes was created by collecting information from previous studies and acknowledged website resources (Arabidopsis Cytochromes P450; Paquette et al., 2000; Ehlting et al., 2008; Nelson, 2009; Bak et al., 2011; Nelson and Werck-Reichhart, 2011; Boutanaev et al., 2015) (http://www.p450.kvl.dk/p450.shtml). Genes marked in Araport 11 as pseudogenes were excluded from the further analysis. Genes were assigned to clans and families according to the information from the above resources. A list of TS genes was created based on a previous study (Boutanaev et al., 2015) and restricted to genes with valid Araport 11 locus. Genotypes were assigned based on criteria defined for RD dataset: (CN =< 1 as losses, CN >=3.4 as gains, the remaining genotypes were classified as unchanged). Genes from thalianol, tirucalladienol, arabidiol/baruol and marneral gene clusters were already genotyped. Gene coordinates were downloaded from Araport 11. All CYP genes positioned at a distance +/- 30 kb from TS gene borders were classified as paired with a given TS gene. Information about predicted secondary metabolism clusters was retrieved from plantiSMASH resource (Kautsar et al., 2017).

### Prediction and analysis of BARS1 and BARS2 3D protein structures

The three-dimensional structures of the reference baruol synthase 1 proteins NP_193272.1, NP_001329547.1, as well as Cvi-0 proteins encoded by *ATCVI-4G38020* (*BARS1*) and *ATCVI-4G38110* (*BARS2*), were predicted from their amino acid sequences using the AlphaFold2 code through the ColabFold software (Jumper et al., 2021; Mirdita et al., 2022). The modeling studies were performed for a single amino acid chain. A crystal structure of human OSC in a complex with lanosterol (ID 1W6K) was retrieved from the Protein Data Bank (Thoma et al., 2004; Berman et al., 2007). The SSM algorithm implemented in COOT was used for superpositions of protein models (Krissinel and Henrick, 2004; Emsley et al., 2010) (**Supplemental information**).

## Supporting information

Supplemental file 3. Superposed 3D models of BARS1, BARS2 and human oxidosqualene cyclase proteins

Supplemental file 4. Gene models for selected genes for Cdm-0, Kn-0 and Ty-1 genomes.

## Data availability

Data generated or analyzed during this study are included in this published article and its supplementary information files. Public sources of previously published WGS, RNA-Seq datasets and catalogs of genetic variants (SNPs and CNVs) are detailed in the Methods section.

## Author contributions

Conceptualization, A.Z.; Methodology, M.M.Z., P.W., and A.Z.; Investigation, M.M.Z., A.S., P.W., P.S., K.B., and T.I.; Software, M.M.Z., A.S., P.W., and M.Z.; Visualization, M.M.Z., K.B., and A.Z.; Formal Analysis – M.M.Z; Writing – Original Draft, M.M.Z., and A.Z.; Writing – Review & Editing, M.M.Z., K.B., M.F., M.Z., and A.Z; Supervision, M.F., and A.Z.; Project Administration, A.Z.; Funding Acquisition, M.F., and A.Z.

## Acknowledgments

We thank Piotr Kozłowski for fruitful discussions and comments on the manuscript. This research was supported in part by PLGrid Infrastructure. The authors declare that they have no competing interests.

## Supplemental information

**Supplemental file 1. Supplemental Tables S1-S17**. Format .xlsx

**Supplemental file 2. Supplemental information and Supplemental Figures S1-S21**. Format .pdf

**Supplemental file 3. Superposed 3D models of BARS1, BARS2 and human oxidosqualene cyclase proteins**. Format .pdb, zipped.

**Supplemental file 4. Gene models for selected genes for Cdm-0, Kn-0 and Ty-1 genomes**. Format .gtf, zipped.

