## Supplemental file 2. Supplemental information and Supplemental Figures S1-S21 for "Copy number variations shape the structural diversity of Arabidopsis metabolic gene clusters and are associated with the climatic gradient"

#### Prediction and analysis of BARS1 and BARS2 3D protein structures

A theoretical 3D model of a plant baruol synthase 1 isoform (NP_193272.1) obtained by the AlphaFold2 algorithm is available (Uniprot ID: O23390). However, an experimental 3D model of any plant OSC is elusive. Thus, we attempted to obtain 3D models of the reference (Col-0) baruol synthase 1 protein (isoforms NP_193272.1, NP_001329547.1) as well as BARS proteins from Cvi-0 encoded by gene duplicates *ATCVI-4G38020* (BARS1) and *ATCVI-4G38110* (BARS2). A comparison of NP_193272.1 models generated with ColabFold software (this study) with that from the UniProt database indicated a very high agreement of their geometrical parameters. Both models superposed with rmsd of ~0.42 Å for 754 Cα atoms, out of overall 759 Cα atoms. Also, the prediction quality for all models we generated with ColabFold was characterized by high pLDDT measures. For the first best predictions, these measures were equal to 91.8 (NP_193272.1), 94.3 (ATCVI-4G38020) and 94.9 (NP_001329547.1 and ATCVI-4G38110). These results indicated that our predictions could be treated with confidence. Therefore, we used 3D protein models obtained with the ColabFold software for further comparative analyses.

Amino acid sequences of Col-0 and Cvi-0 homologs were very similar. Not surprisingly, their structural models superimposed each other very well, with rmsd values from 0.26 Å to 0.42 Å for the Cα atoms (**Supplemental Table S10**). On the other hand, a simple structural comparison of these plant enzymes did not provide any information about their active site. Therefore, we surveyed the Protein Data Bank in search of any OSC experimental structure determined in a complex with its ligand. Using an amino acid sequence of the Col-0 baruol synthase 1 enzyme NP_193272.1, we identified a crystal structure of human OSC (~35% sequence identity) in a complex with lanosterol (ID 1W6K). Superposition of our 3D models of BARS homologs with human OSC revealed their structural similarity, with rmsd values around 1.2 Å for Cα atoms. Moreover, the presence of lanosterol molecule in the active site of the human enzyme allowed us to identify potential substrate-binding cavities in plant homologs. It is of note that the catalytic aspartate residue D455 present in human cyclase had its counterparts in plant BARS homologs (D493 in Col_BARS1 NP_193272.1, D490 in NP_001329547.1 as well as in ATCVI-4G38020 and ATCVI-4G38110).

### Protein sequences of genes predicted with Augustus in de novo genomic assemblies

##### Ecotype Mir-0

Marneral gene cluster

>g1_Mir0 96 aa; Best reference protein match NP_199072.1 AT5G42580 499 aa
MFKPERFLVSSISGDEEKIREQAVKYVTFGGGRRTCPAVKLAHIFMETAIGAMVQCFDWR
IKGEKVYMEEAVSGLSLKMAHPLKCTPVVRFDPFSF

##### Ecotype Mitterberg-2-185

Thalianol gene cluster

>g1_Mitterberg 749 aa; Best reference protein match NP_199605.1 AT5G47940 749 aa
MGASNHDNDFNSTTNWKLVDGTLIDAISFESSFTANPESDDGIISAAVDHVTKSPLLLLP
PVPNGEPCEITITFAQEHELRQIYIRSSARVYEVYYTKKRRHDKEYLCTVRCGVAIRDEE
VLQIPLTESADSKPVKDLIERKVTDNGNGRTSEDDWVEVKASDDSLLNNEKQDFYEATAE
INDAEPCTSITVRLLSLQDKRCALVDEVYVFADPVDPSESEKEEATGTGNSSSSSLMAMF
MPALLQLSRGKDVRKERDIQVSDKSNSTDPVAIGNTDQIGVSSPVLVDTVAKQVDAATRV
SGEESKPSISCNNVETIMDQLVKKVSMIETILIRFEDQMLKPINSIDARLQLVEKKLEQL
GNKSFESDLGFRKKIPNQDSLRSDTDKTPDTDESDGLTKNTDVVPDSSSIDNSEDCAVVL
PKNRLDNILSKSVELESENSSISGNEMISAEPEISNEEVGHSFEEKPKYSLSINDALASA
LAGLLSSHSITDGKYSQALVITAPEFSSEDDVEIEQKPGTSAHPDDSQVAAEESENRYSS
SLESSTSSQKEPGITPDDSHGTMYGVFKKLDDSFGGDEEAETVVSVSDNALDEEMVTSST
KADCYTERKNLSYKPTEPDSLIHELESSNVTTAKCKGEPSMDDVLKSVLGFQPTTSSVDF
LTPVLDVKFNLENKDSDSKYFFEVLFTGESKTYLDCKNDVFDDNLVSVEDEEELKGPPTD
TLSSVEMNHYATNEMPIHWNGEISEASLI

>g2_Mitterberg 387 aa; Best reference protein match NP_001078731.1 AT5G47970 387 aa
MTVSEAYSPPLFSIAPMMGWTDNHYRTLARLITKHAWLYTEMLAAETIVYQEDNLDSFLA
FSPDQHPIVLQIGGRNLENLAKATRLANAYAYDEINFNCGCPSPKVSGRGCFGALLMLDP
KFVGEAMSVIAANTNAAVTVKCRIGVDDHDSYNELCDFIHIVSSLSPTKHFIIHSRKALL
SGLSPSDNRRIPPLKYEFFFALLRDFPYLKFTINGGINSVVEADAALRSGAHGVMLGRAV
YYNPWHILGHVDTVIYGSPSSGITRRQVLEKYKVYGESVLGKYGKGRPNLRDIVRPLINL
FHSESGNGQWKRRTDAALLHCTTLQSFLDEVLPAIPDYVLDSSAVKEATGREDLFADVQR
LLPPPYEKESLKALERMPTRPVILDEE

>g3_Mitterberg 223 aa; Best reference protein match NP_199607.1 AT5G47960 223 aa
MSKFQSNFNQKIDYVFKVVLIGDSAVGKSQLLARFSRNEFSIESKATIGVEFQTRTLEID
RKTIKAQIWDTAGQERYRAVTSAYYRGAVGAMLVYDITKRQSFDHVARWLEELRGHADKN
IVIMLIGNKTDLGTLRAVPTEDAKEFAQRENLFFMETSALDSNNVEPSFLTVLTEIYRIV
SKKNLVANEEGESGGDSSLLQGTKIVVAGEETESKGKGCCGTS

>g4_Mitterberg 426 aa; Best reference protein match NP_199606.1 AT5G47950 426 aa
MDTMKVETIGKEIIKPSATTPNDLPTLQLSIMDILMPPVYAVAFLFYTKDDLISQEQTSH
TLKTSLSEILTKFHPLAGRVNGVTIKSTDEGAVFVEARVDNCDLSGFLRSPDTESLKQLL
PVDDEPAPTWPLLVVKATYFQCGGMAIGLCISHRLADAASLSIFLQAWAATARGESDSVA
SPEFCSTKLYPAANEAIKIPGEVVKRTSVTKRFVFVASKIEELRKKVASDVVPRPTRVQS
VTSLIWKCAVTASTDKIREKALFQPANLRTKIPSLLSENQIGNFLFNSLTLDGKAGVDIV
ETVKELQKRAEELSGLVQHEEGSSMTIGSRLFGEIINSKFNFELHDMHSVTSWCKIPLYD
ACFGWGSPVWVAGSVSPDLENVTVLIDSKDGQGIEAWVTLHQDNMLLFEQSTELLAFASP

NPSVLI

>g5_Mitterberg 404 aa; Best reference protein match NP_199609.1 AT5G47980 443 aa
MIFFYNLADLAEKSPDIVSTRLRSSLSQALSRFYPLAGKKEGVSISCNDEGAVFTEARTN
LLLSDFLRNIDINSLKILIPTLAPGESLDSRPLLSVQATFFGSGSGVAVEICVSHCICDA
ASVSTFFRGWAATARGDSNDELSTPQFAEVAIHPPADISIHGSPFNALSEVREKCVTNRF
VFESDKITKLKIVAASKSVPSPTRVEAVMSLIWRCARNASHANLIVPRATMMTQSMDLRL
RIPTNVLSPDAIGNLQGVFFLKRGPGSEIEISEVVAEFRKEKEEFNEMIKENVNGGHTNT
TLGQKIMSGIANYMSELKPNIDTYTMSSWCRKAFYEVDFGWGRPAWVGLGHQDIQDGVMY
VLLVDAKDGEGVEAWVGIPEQDMAAFVCDQELLSYASLNPPVLI

>g6_Mitterberg 404 aa; Best reference protein match NP_199609.1 AT5G47980 443 aa
MIFFYNLADLAEKSPDIVSTRLRSSLSQALSRFYPLAGKKEGVSISCNDEGAVFTEARTN
LLLSDFLRNIDINSLKILIPTLAPGESLDSRPLLSVQATFFGSGSGVAVEICVSHCICDA
ASVSTFFRGWAATARGDSNDELSTPQFAEVAIHPPADISIHGSPFNALSEVREKCVTNRF
VFESDKITKLKIVAASKSVPSPTRVEAVMSLIWRCARNASHANLIVPRATMMTQSMDLRL
RIPTNVLSPDAIGNLQGVFFLKRGPGSEIEISEVVAEFRKEKEEFNEMIKENVNGGHTNT
TLGQKIMSGIANYMSELKPNIDTYTMSSWCRKAFYEVDFGWGRPAWVGLGHQDIQDGVMY
VLLVDAKDGEGVEAWVGIPEQDMAAFVCDQELLSYASLNPPVLI

>g7_Mitterberg 511 aa; Best reference protein match NP_199610.1 AT5G47990 511 aa
MASMITVDFENCFIFLLLCLFSRLSYDLFFRKTKDLRAGCALPPSPPSLPIIGHLHLILF
VPIHQSFKNISSKYGPLLHLRFFNFPIVLVSSASTAYEIFKAQDVNVSSRPPPPIEESLI
LGSSSFINTPYGDYSKFMKKFMVQKLLGPQALQRSRNIRADELERFYKTLLDKAMKKQTV
EIRNEAMKLTNNTICKMIMGRSCSEENGEAETVRGLVTESIFLTKKHFLGAMFHKPLKKL
GISLFAKELMNVSNRFDELLEKILVEHEEKLQEHHQTSDMLDMLLEAYGDENAEYKITRD
QIKSLFVDLFSAGTEASANTIQWTMAEIIKNPKICERLREEIDSVVGKTRLVQETDLPNL
PYLQAIVKEGLRLHPPGPVVRTFKETCEIKGFYIPEKTRLFVNVYAIMRDPDFWEDPEEF
KPERFLASSRLGEEDEKREDMLKYIPFGSGRRACPGSHLAYTVVGSVIGMMVQHFDWIIK
GEKINMKEGGTMTLTMAHPLKCTPVPRNLNT

>g8_Mitterberg 471 aa; Best reference protein match NP_001032030.1 AT5G48000 477 aa
MSFVWSAAVWVIAVAAVVISKWLYRWSNPKCNGKLPPGSMGLPIIGETCDFFEPHGLYEI
SPFVKKRMLKYGPLFRTNIFGSNTVVLTEPDIIFEVFRQENKSFVFSYPEAFVKPFGKEN
VFLKHGNIHKHVKQISLQHLGSEALKKKMIGEIDRVTYEHLRSKANQGSFDAKEAVESVI
MAHLTPKIISNLKPETQATLVDNIMALGSEWFQSPLKLTTLISIYKVFIARRDALQVIKD
VFTRRKASREMCGDFLDTMVEEGEKEDVIFNEESAINLIFAILVVAKESTSSVTSLAIKF
LAENHKALAELKREHAAILQNRNGKGAGVSWEEYRHQMTFTNMKGCERCRNQRQVSLKHG
STRSYTIPAGWIVAVIPPAVHFNDAIYENPLEFNPWRWEGKELRSGSKTFMVFGGGVRQC
VGAEFARLQISIFIHHLVTTYDFSLAQESEFIRAPLPYFPKGLPIKISQSL

>g9_Mitterberg 764 aa; Best reference protein match NP_001078733.1 AT5G48010 766 aa
MWRLRTGPKAGEDTHLFTTNNYAGRQIWEFDANAGSPQEIAEVEDARHKFSDNTSRFKTT
ADLLWRMQFLREKKFEQKIPRVIIEDARKIKYEDAKKALKRGLLYFTALQADDGHWPAEN
SGPNFYTPPFLICLYITGHLEKIFTPEHVKELLRHIYNMQNEDGGWGLHVESHSVMFCTV
INYVCLRIVGEEVGHDDQRNGCAKAHKWIMDHGGATYTPLIGKALLSVLGVYDWSGCNPI
PPEFWLLPSSFPVNGGTLWIYLRDTFMGLSYLYGKKFVATPTPLILQLREELYPEPYAKI
NWTQTRNRCGKEDLYYPRSFLQDLFWKSVHMFSESILDRWPLNKLIRQRALQSTMALIHY
HDESTRYITGGCLPKAFHMLACWIEDPKSDYFKKHLARVREYIWIGEDGLKIQSFGSQLW
DTALSLHALLDGIDDHDVDDEIKTTLVKGYDYLKKSQITENPRGDHFKMFRHKTKGGWTF
SDQDQGWPVSDCTAESLECCLFFESMPSELIEKKMDVEKLYDAVDYLLYLQSDNGGIAAW
QPVEGKAWLEWLSPVEFLEDTIVEYVECTGSAIAALTQFNKQFPGYKNVEVKRFITKAAK

YIEDMQTVDGSWYGNWGVCFIYGTFFAVRGLVAAGKTYSNCEAIRKAVRFLLDTQNTEGG
WGESFLSCPSKKYTPLKGNSTNVVQTAQALMVLIMGDQMERDPLPVHRAAQVLINSQLDN
GDFPQQEIMGTFMRTVMLHFPTYRNTFSLWALTHYTHALRRLLP

>g10_Mitterberg 355 aa; Best reference protein match NP_568689.1; AT5G48020 355 aa
MELPVVDLSRYLDFSGDELGSDLLESCRQVSRILKETGALIVKDPRCCAQDNDRFIDMME
NYFEKPDDFKRLQQRPNLHYQVGATPEGVEVPRSLVDEEMQEKFKTMPNEYKPHIPKGPD
HKWRYMWRVGPRPSNTRFKELNSEPVIPEGFPEWEEVMDSWGFKMISAVEVVAEMAAIGF
GLPKDAFTSLMKQGPHLLAPTGSDLNCYNEEGTIFAGYHYDLNFLTIHGRSRFPGLYIWL
RNGEKVAVKVPVGCLLIQAGKQIEWLTAGECIAGMHEVVVTSKTKDAITLAKEQNRSLWR
VSSTLFAHIASDAELKPLGHFAESSLASKYPAIPAGEYVEQELSVINLKGNKGFS

**Ecotype Dolna-1-40**

Tirucalladienol gene cluster

>g1_Dolna 318 aa; Best reference protein match NP_198463.1 AT5G36140 318 aa
MYLTIIFLFISSIIFPLLFFLGKHLSNFRYPNLPPGKIGFPLIGETLSFLSAGRQGHPEK
FVTDRVRHFSSGIFKTHLFGSPFAVVTGASGNKFLFTNENKLVISWWPDSVNKIFPSSTQ
TSSKEEAIKTRMLLMPSMKPEALRRYVGVMDEIAQKHFETEWANQDQLIVFPLTKKFTFS
IACRLFLSMDDLERVRKLEEPFTTVMTGVFSIPIDLPGTRFNRAIKASRLLSKEVSTIIR
QRKEELKAGKVSVEQDILSHMLMNIGETKDEDLADKIIALLIGGHDTTSIVCTFVVNYLA
EFPHIYQRVLEGMQIPLL

**Ecotype Cvi-0**

Arabidiol/baruol gene cluster

>g1_Cvi 513 aa; Best reference protein match NP_193270.1 AT4G15350 509 aa; Augustus predicted gene (chr4:8591612-8593232 complement)
MAAMIFILLCLFTFLCYSLFYKKPKDSRANCDRPPSPPSLPIIGHLHLILSNLAHKSFQR
LSSKYGPLLHLRIFHIPIVLVSSASVAYDIFRAQDVNVSFRSTSTFEECLFFGTSGFFQA
PYGDYWKFMRKLMVTKLLGPQALERSRNIRVEEIDRLYKNLLNKAMKKESVEIGEEASKL
SNNVICTMIMGRSCSEDNGEAERMRSLVSEAMALTKKFFLANIFHKPLKMLGISLFEKEI
MSVSHKFDELLEKILVEHEEKMEEHHQGTDMMDVLLEAYRDENATYKITRNQIKSLIVEL
LIAGTDTSATTTQWIMAELINHPKVFERVREEIDLVVGRSRLIQETDLPNLAYLQAVVKE
ALRLHPPGPLVPRTLQESCEIKGYYIPEKTIVIVNSYAVMRDPYVWEDPEEFKPERFLDI
SSSVQEEEISDKILKFIPFASGRRGCPGTNLAYINVETAIGVMVQCFDWIIKGKEVNMSE
AAGTMVLTLAEPLMCTPVARTLNPLPASLRAYS

**Ecotype Eri-1**

Arabidiol/baruol gene cluster

>g1_Eri 763 aa; Best reference protein match NP_001329547.1 AT4G15370 756 aa; Augustus predicted gene (chr4:8744883-8749727 complement)
MWRLRIGAKAKDNTHLFTTNNYVGRQIWEFDANAGSPEELAEVEEARRNFSNNRSRFKAS
ADLLWRMQFLREKKFEQKIPRVIVEDAEKITYEDAKTALRRGILYFTALQADDGHWPAEN
AGSIFFNAPFVICLYITGHLEKIFTHEHRVELLRYMYNHQNEDGGWGLHVESPSNMFCSV
INYICLRILGVEAGHDDKGSACARARKWILDHGGATYSPLIGKAWLSVLGVYDWSGCKPI
PPEFWFLPSFFPVNGGTLWIYLRDIFMGLSYLYGKNFVATSTPLILQLREEIYPDPYTNI
SWRQARNRCAKEDLYYPQSFLQDLFWKGVHVFSENILNRWPFNNLIRQRALRTTMELVHY
HDEATRYITGGSVPKVFHMLACWVEDPESDYFKKHLARVPDFIWIGEDGLKIQSFGSQVW
DTALSLHVFIDGFDDDVDEEIRSTLLKGYDYLEKSQVTENPPGDYMKMFRHMAKGGWTFS
DQDQGWPVSDCTAESLECCLFFESMSSEFIGKKMDVEKLYDAVDFLLYLQSDNGGITAWQ

PADGKTWLEWLSPVEFIEDAVVEHEYVECTGSAIVALAQFNKQFPGYKKEEVERFITKGV
KYIEDLQMVDGSWYGNWGVCFIYGTFFAVRGLVAAGKCYNNCEAIRRAVRFILDTQNTEG
GWGESYLSCPRKKYIPLIGNKTNVVNTGQALMVLIMGNQMKRDPLPVHRAAKVLINSQMD
NGDFPQQEIMGVFKMNVMLHFPTYRNMFTLWALTHYTKALRGL

>g2_Eri 513 aa; Best reference protein match NP_193270.1 AT4G15350 509 aa;
Augustus predicted gene (chr4:8792845-8794465 complement)
MAAMIFILLCLFTFLCYSLFYKKPKDSRANCDRPPSPPSLPIIGHLHLILSNLAHKSFQR
LSSKYGPLLHLRIFHIPIVLVSSASVAYDIFRAQDVNVSFRSTSTFEECLFFGTSGFFQA
PYGDYWKFMRKLMVTKLLGPQALERSRNIRVEEIDRLYKNLLNKAMKKESVEIGEEASKL
SNNVICTMIMGRSCSEDNGEAERMRSLVSEAMALTKKFFLANIFHKPLKMLGISLFEKEI
MSVSHKFDELLEKILVEHEEKMEEHHQGTDMMDVLLEAYRDENATYKITRNQIKSLIVEL
LIAGTDTSATTTQWIMAELINHPKVFERVREEIDLVVGRSRLIQETDLPNLAYLQAVVKE
ALRLHPPGPLVPRTLQESCEIKGYYIPEKTIVIVNSYAVMRDPYVWEDPEEFKPERFLDI
SSSVQEEEISDKILKFIPFASGRRGCPGTNLAYINVETAIGVMVQCFDWIIKGKEVNMSE
AAGTMVLTLAEPLMCTPVARTLNPLPASLRAYS

**Ecotype Ler-0**

Arabidiol/baruol gene cluster

>g1_Ler 763 aa; Best reference protein match NP_001329547.1 AT4G15370 756 aa;
Augustus predicted gene (chr4:9215522-9220364 complement)
MWRLRIGAKAKDNTHLFTTNNYVGRQIWEFDANAGSPEELAEVEEARRNFSNNRSRFKAS
ADLLWRMQFLREKKFEQKIPRVIVEDAEKITYEDAKTALRRGILYFTALQADDGHWPAEN
AGSIFFNAPFVICLYITGHLEKIFTHEHRVELLRYMYNHQNEDGGWGLHVESPSNMFCSV
INYICLRILGVEAGHDDKGSACARARKWILDHGGATYSPLIGKAWLSVLGVYDWSGCKPI
PPEFWFLPSFFPVNGGTLWIYLRDIFMGLSYLYGKNFVATSTPLILQLREEIYPDPYTNI
SWRQARNRCAKEDLYYPQSFLQDLFWKGVHVFSENILNRWPFNNLIRQRALRTTMELVHY
HDEATRYITGGSVPKVFHMLACWVEDPESDYFKKHLARVPDFIWIGEDGLKIQSFGSQVW
DTALSLHVFIDGFDDDVDEEIRSTLLKGYDYLEKSQVTENPPGDYMKMFRHMAKGGWTFS
DQDQGWPVSDCTAESLECCLFFESMSSEFIGKKMDVEKLYDAVDFLLYLQSDNGGITAWQ
PADGKTWLEWLSPVEFIEDAVVEHEYVECTGSAIVALAQFNKQFPGYKKEEVERFITKGV
KYIEDLQMVDGSWYGNWGVCFIYGTFFAVRGLVAAGKCYNNCEAIRRAVRFILDTQNTEG
GWGESYLSCPRKKYIPLIGNKTNVVNTGQALMVLIMGNQMKRDPLPVHRAAKVLINSQMD
NGDFPQQEIMGVFKMNVMLHFPTYRNMFTLWALTHYTKALRGL

>g2_Ler 513 aa; Best reference protein match NP_193270.1 AT4G15350 509 aa;
Augustus predicted gene (chr4:9263484-9265104 complement)
MAAMIFILLCLFTFLCYSLFYKKPKDSRANCDRPPSPPSLPIIGHLHLILSNLAHKSFQR
LSSKYGPLLHLRIFHIPIVLVSSASVAYDIFRAQDVNVSFRSTSTFEECLFFGTSGFFQA
PYGDYWKFMRKLMVTKLLGPQALERSRNIRVEEIDRLYKNLLNKAMKKESVEIGEEASKL
SNNVICTMIMGRSCSEDNGEAERMRSLVSEAMALTKKFFLANIFHKPLKMLGISLFEKEI
MSVSHKFDELLEKILVEHEEKMEEHHQGTDMMDVLLEAYRDENATYKITRNQIKSLIVEL
LIAGTDTSATTTQWIMAELINHPKVFERVREEIDLVVGRSRLIQETDLPNLAYLQAVVKE
ALRLHPPGPLVPRTLQESCEIKGYYIPEKTIVIVNSYAVMRDPYVWEDPEEFKPERFLDI
SSSVQEEEISDKILKFIPFASGRRGCPGTNLAYINVETAIGVMVQCFDWIIKGKEVNMSE
AAGTMVLTLAEPLMCTPVARTLNPLPASLRAYS

**Ecotype C24**

Arabidiol/baruol gene cluster

>g1_C24 78 aa; Best reference protein match NP_001329547.1 AT4G15370 756 aa;
Augustus predicted gene (chr4:9580438-9581743 complement)

MWKLIIGSKAGDDIHLFSTNNYVGRQIWEFDAKAGSPEELAEVEEARQNFTDNRSHFKAS

```
ADLLWRMQFLREKKFEQKIPRVIIEDAEKITYEDAKTALKRGLLYFTALQADDGHWPAEN
AGSIFFNAPFVICMYITGHLERIFTPEHVRELLRYLYNHQNEDGGWGLHIESPSNMFCTV
INYICLRILGVEAGYDDEGSACARARKWILDHGGATYSPLIGKAWLSVLGVYDWSGCKPI
PPEFWLLPSFLPVNGGLKLEGL
```

**Distribution of accessions with varying CYP705A2-BARS1 copy number status**

Map available at:
https://www.google.com/maps/d/edit?mid=1ZaAMX-EDYlbBtKbKdBsS06HMjvc&usp=sharing
Last accessed: September 21, 2022.

**Supplemental Figures**

Figure S1. IGV screens of genomic regions covering Arabidopsis MGCs

Figure S2. Copy number analysis of genes in thalianol (A), marneral (B) and tirucalladienol (C) gene clusters

Figure S3. Copy number analysis of genes in arabidiol/baruol gene cluster

Figure S4. Evidence supporting manual correction of genotype assignments in individual genes and accessions

Figure S5. WGS data-based evidence for a new type of deletion in the thalianol gene cluster spanning *CYP705A5*, *CYP708A2* and *THAS1*

Figure S6. Duplication of acyltransferase gene in Mitterberg-2-185

Figure S7. Partial deletion of *CYP705A12* in Mir-0

Figure S8. Differences between the countries in read coverage and mapping indicate that structural variants in tirucalladienol cluster genes are of local origin

Figure S9. Alternative *CYP716A2* gene models

Figure S10. Variation in WGS data coverage and mapping in the region spanning *CYP705A2*, *CYP705A3* and *BARS1* genes

Figure S11. *CYP705A2* duplication detected by RD assay correlates with the occurrence of pseudo-heterozygous SNPs in *CYP705A2* and *BARS1* loci

Figure S12. Multiple sequence alignment of *BARS1* genomic sequences reveals a common lack of the largest intron

Figure S13. Comparison of baruol synthase 1 protein NP_001329547.1 with proteins encoded by *BARS2* genes in Cvi-0, Eri-1 and Ler-0

Figure S14. Heterozygous SNPs in Cvi-0 co-localize with sequence differences between BARS1and its duplicate

Figure S15. Sequence comparison of *CYP705A2* and its duplicate *CYP705A2a*

Figure S16. PCR verification of group assignments based on the presence/absence of *CYP705A2*, *BARS1*, *CYP705A2a* and *BARS2* genes

Figure S17. Spread of PP-AA and PP-PP variants of arabidiol/baruol gene cluster in Arabidopsis population

Figure S18. Latitudes of origin among accessions with and without *CYP705A2a-BARS2* genes divided by country

Figure S19. The combined effect of thalianol and arabidiol/baruol gene clusters' structural variation on root growth phenotypic variation

Figure S20. Expression of selected genes from arabidiol/baruol gene cluster in leaves in accessions from PP-AA and PP-PP groups

Figure S21. Copy number variation of TS/CYP gene pairs in Arabidopsis genome

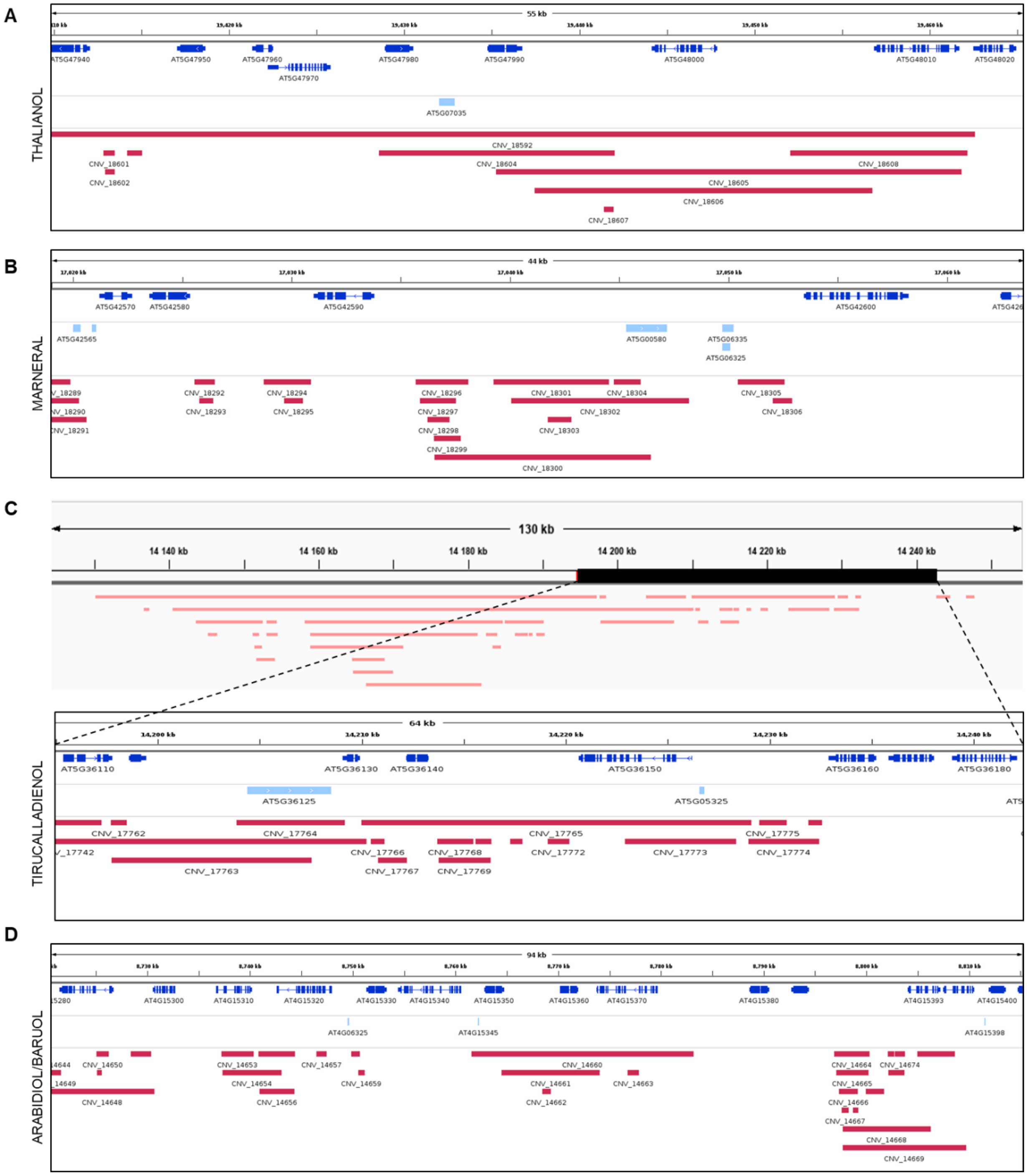


**Figure S1. IGV screens of genomic regions covering Arabidopsis MGCs.** A) Thalianol gene cluster; B) Marneral gene cluster; C) Tirucalladienol gene cluster; D) Arabidiol/baruol gene cluster. Dark blue – protein-coding genes. Araport 11 genomic loci are denoted. Light blue – noncoding genes. Red – CNVs extracted from AthCNV atlas (http://athcnv.ibch.poznan.pl/).

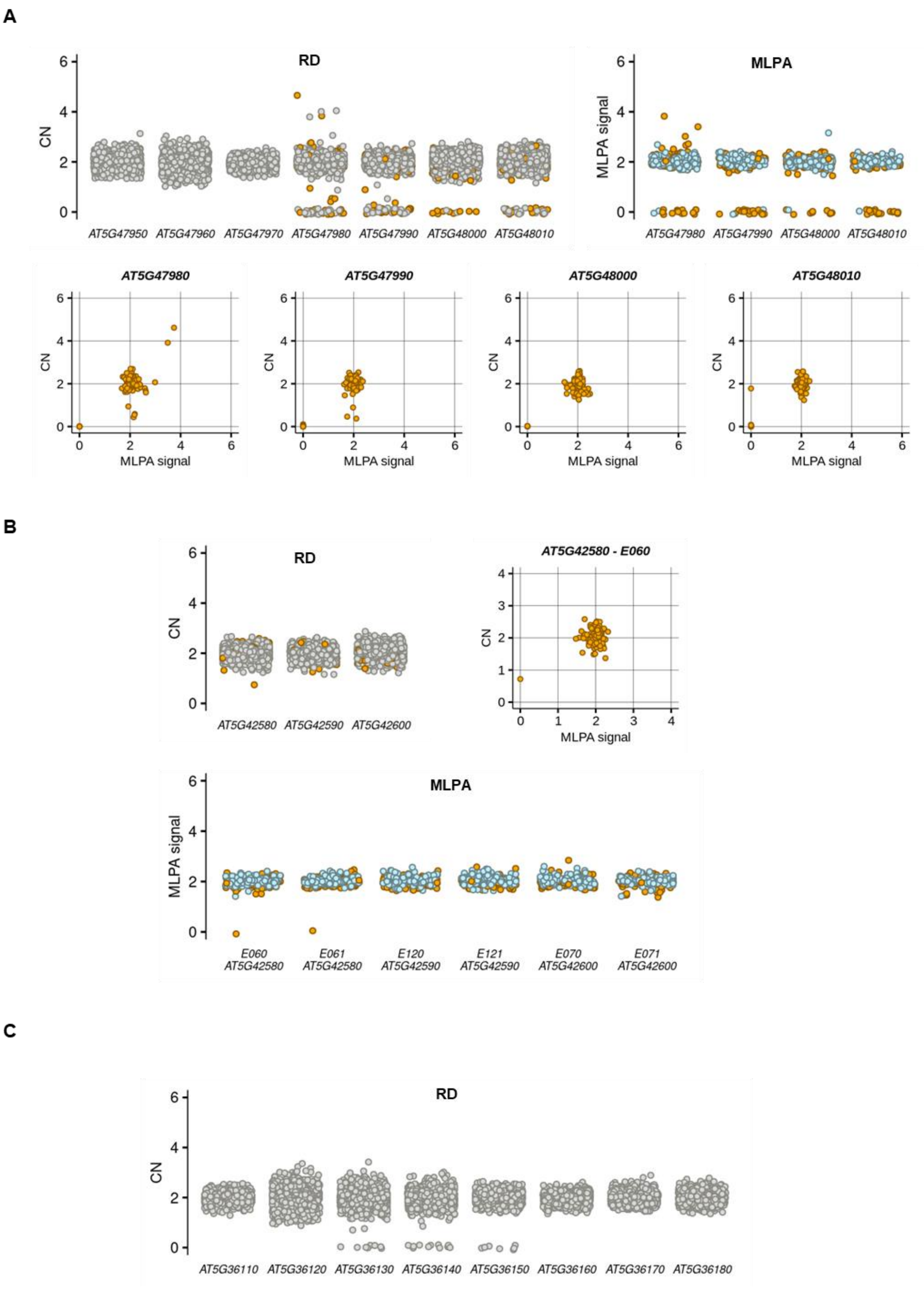


**Figure S2. Copy number analysis of genes in thalianol (A), marneral (B) and tirucalladienol (C) gene clusters.** Dots indicate individual accessions. Colours indicate a method of analysis: grey – RD only, blue – MLPA only, orange – both. The sets of analysed accessions differ between the methods (1,056 for RD; 232 for MLPA) but are identical for each gene.

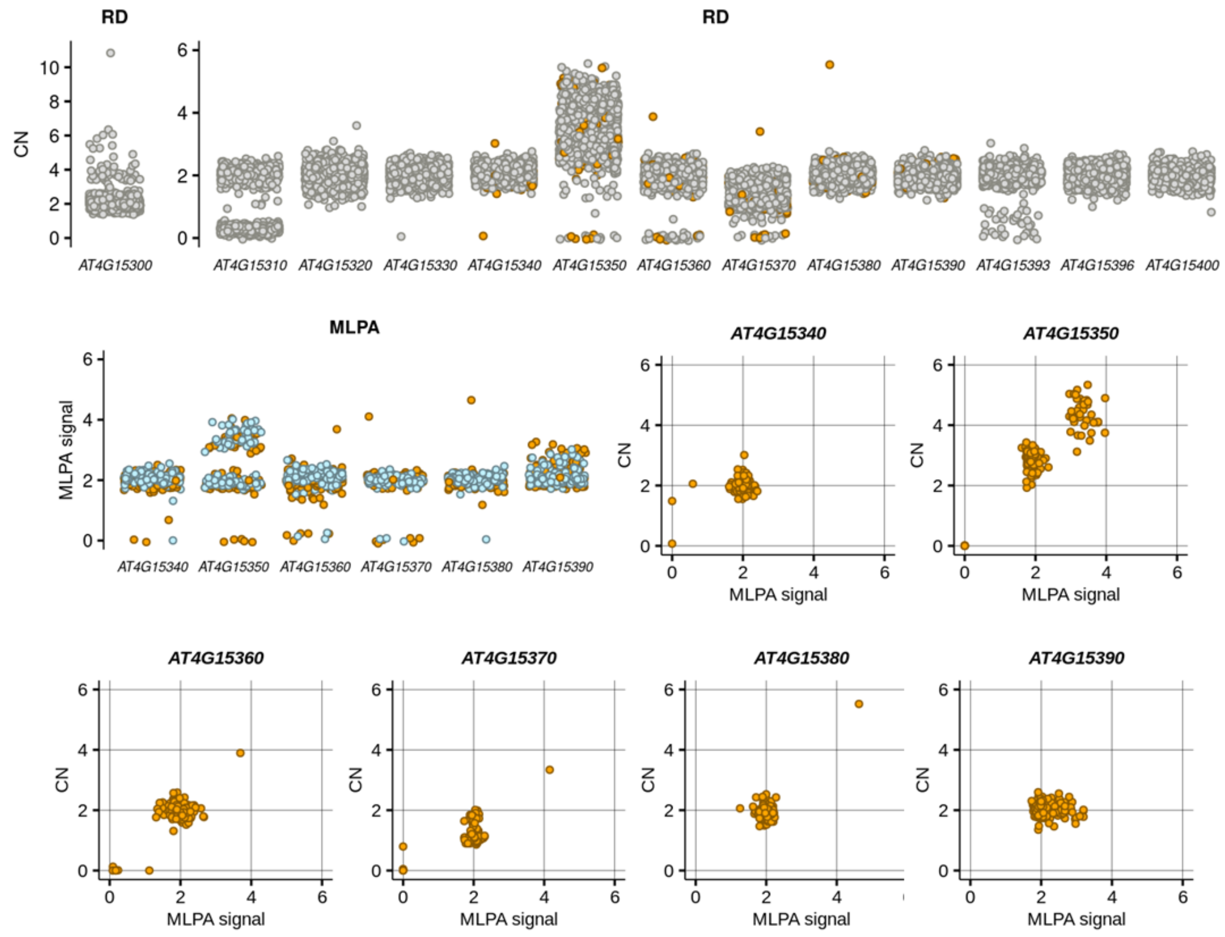


**Figure S3. Copy number analysis of genes in arabidiol/baruol gene cluster.** Dots indicate individual accessions. Colours indicate a method of analysis: grey – RD only, blue – MLPA only, orange – both. The sets of analysed accessions differ between the methods (1,056 for RD; 232 for MLPA) but are identical for each gene.

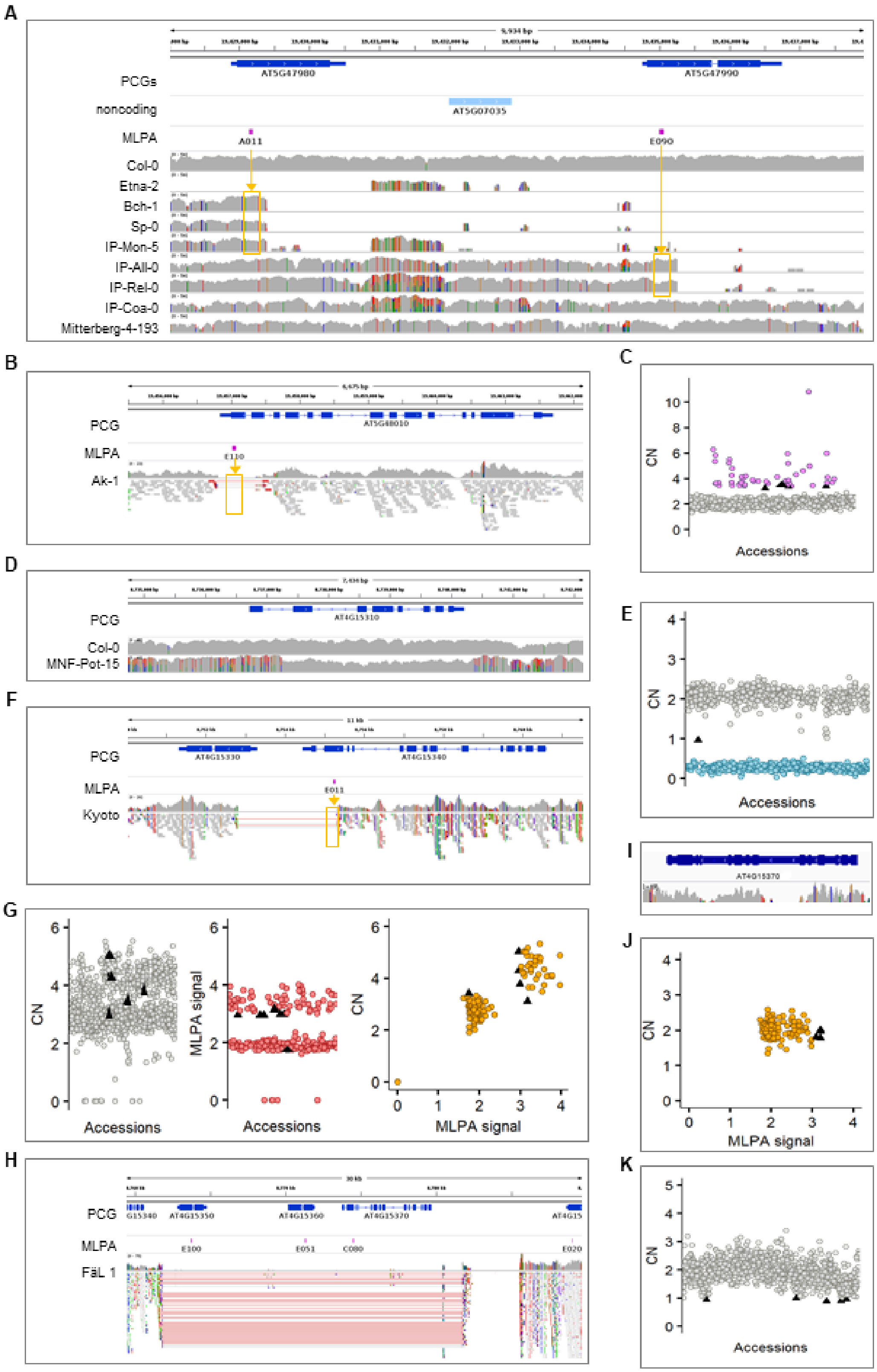


**Figure S4. Evidence supporting manual correction of genotype assignments in individual genes and accessions.** Details are presented in Supplemental Table S7.

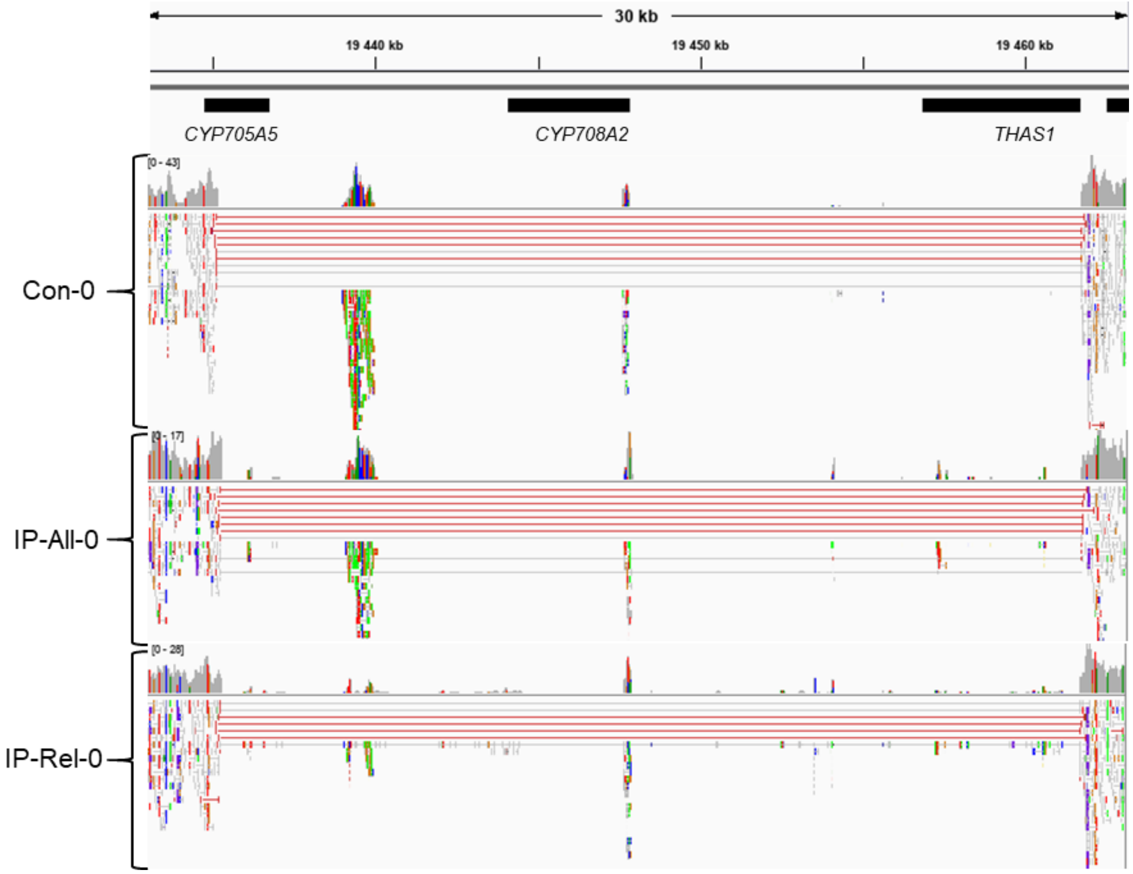


**Figure S5. WGS data-based evidence for a new type of deletion in the thalianol gene cluster spanning *CYP705A5*, *CYP708A2* and *THAS1*.**

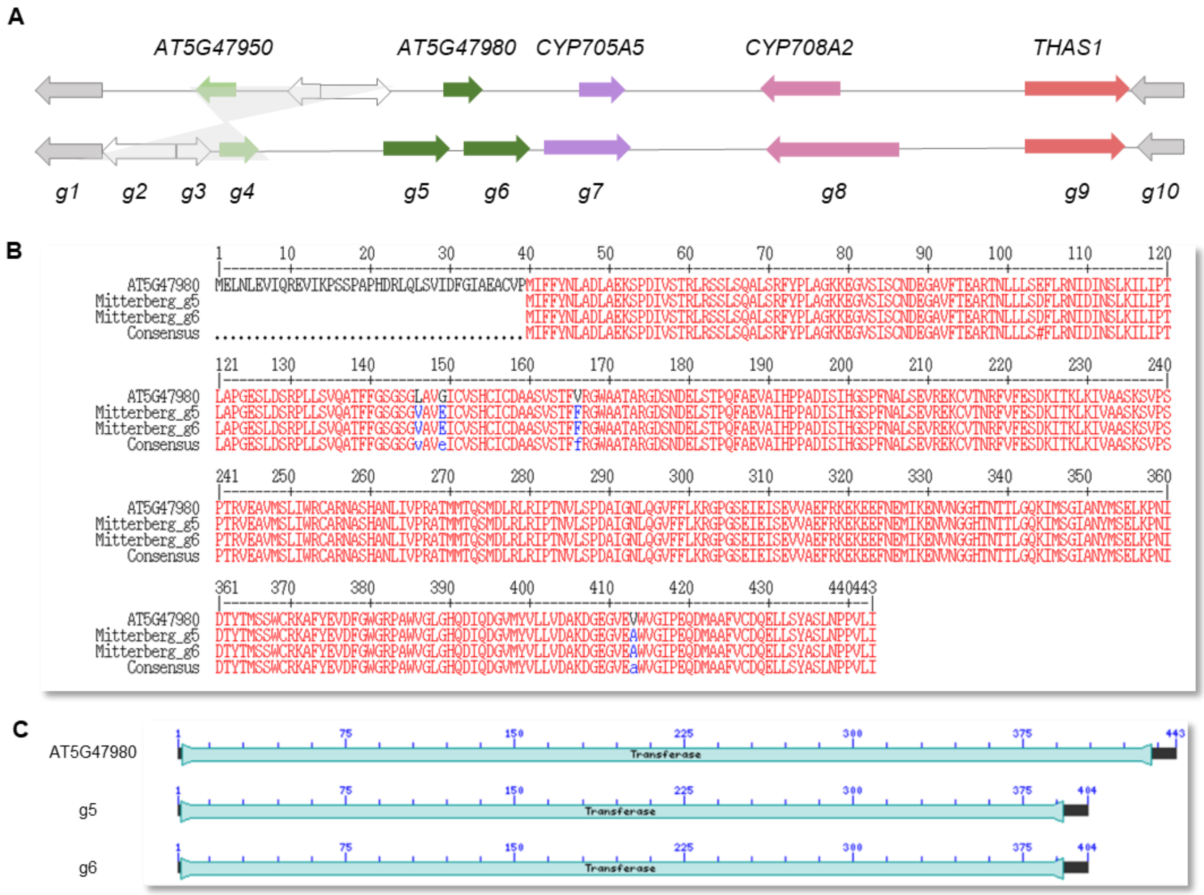


**Figure S6. Duplication of acyltransferase gene in Mitterberg-2-185.** A) Thalianol gene cluster organization in Col-0 (upper) and Mitterberg-2-185 *de novo* assembly (lower). Corresponding genes have the same colors. The region of inversion is marked in grey. B) Sequence alignment of reference AT5G47980 protein with predicted proteins from Mitterberg-2-185. C) Conserved domain prediction. Transferase – transferase domain (pfam02458).

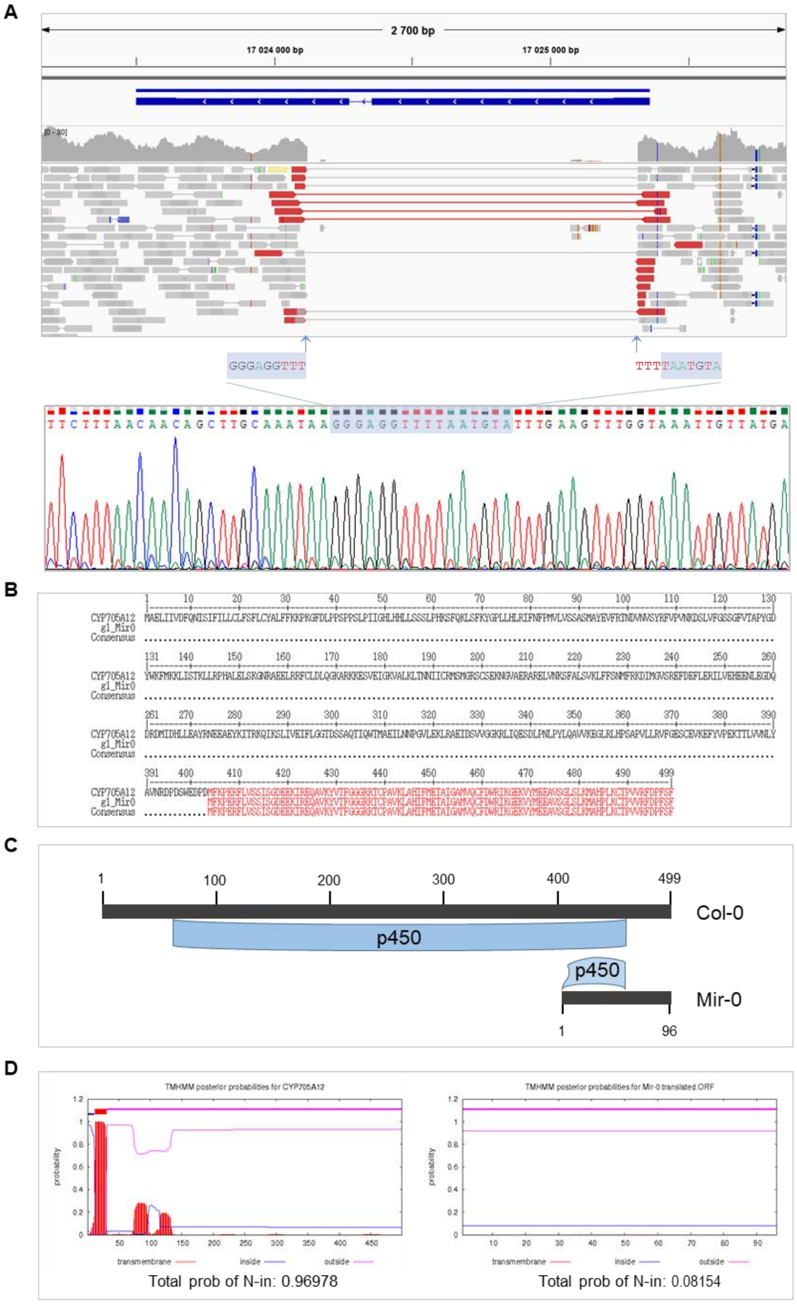


**Figure S7. Partial deletion of *CYP705A12* in Mir-0.** A) WGS-based and Sanger sequencing-based data concordantly indicate there is a 1,202-bp deletion in *CYP705A12* gene sequence in Mir-0. The reference *CYP705A12* model is presented in blue. B) Predicted truncated protein in Mir-0. Multiple protein alignment made with Multalin. C) Full (reference) and partial (Mir-0) cytochrome P450 superfamily domain present in the respective proteins. D) TMHMM server prediction probabilities for CYP705A12 (left) and its truncated homolog in Mir-0 (right).

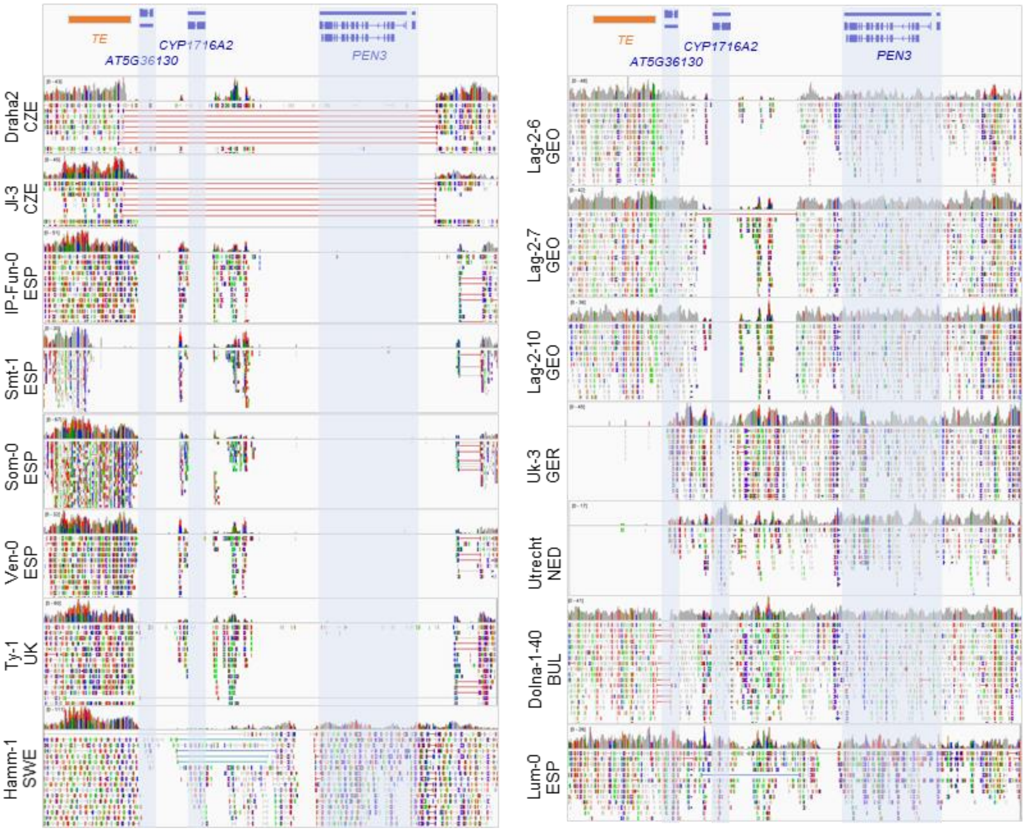


**Figure S8. Differences between the countries in read coverage and mapping indicate that structural variants in tirucalladienol cluster genes are of local origin**. The picture presents data for all 15 accessions with detected copy number changes in tirucalladienol cluster genes.

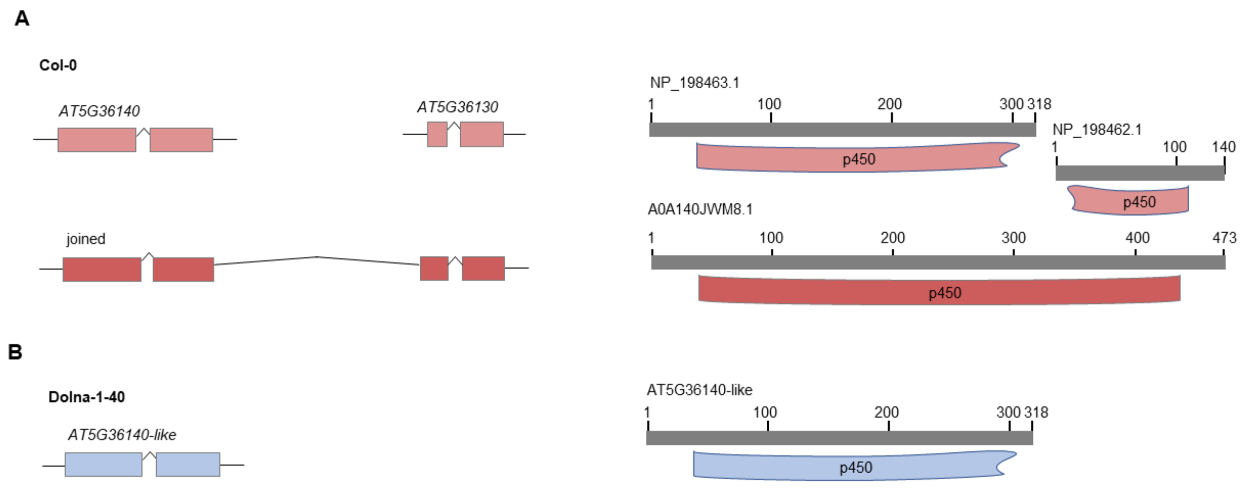


**Figure S9. Alternative *CYP716A2* gene models**. A) According to Araport 11 annotation, there are two separate genes, *AT5G36130* and *AT5G36140* (*CYP716A2*) in the genome. The Augustus tool predicted the same gene models and additionally an alternative joint model. The joint gene encodes a protein identical to the one predicted by Yatsumoto et al. (2016), for which full-length cDNA had been isolated. B) In Dolna-1-40 *AT5G36130* is absent, as indicated by the analysis of its *de novo* genomic assembly. The predicted ORF encodes a protein identical to *AT5G36140*, which lacks C-part of p450 superfamily domain (cl12078).

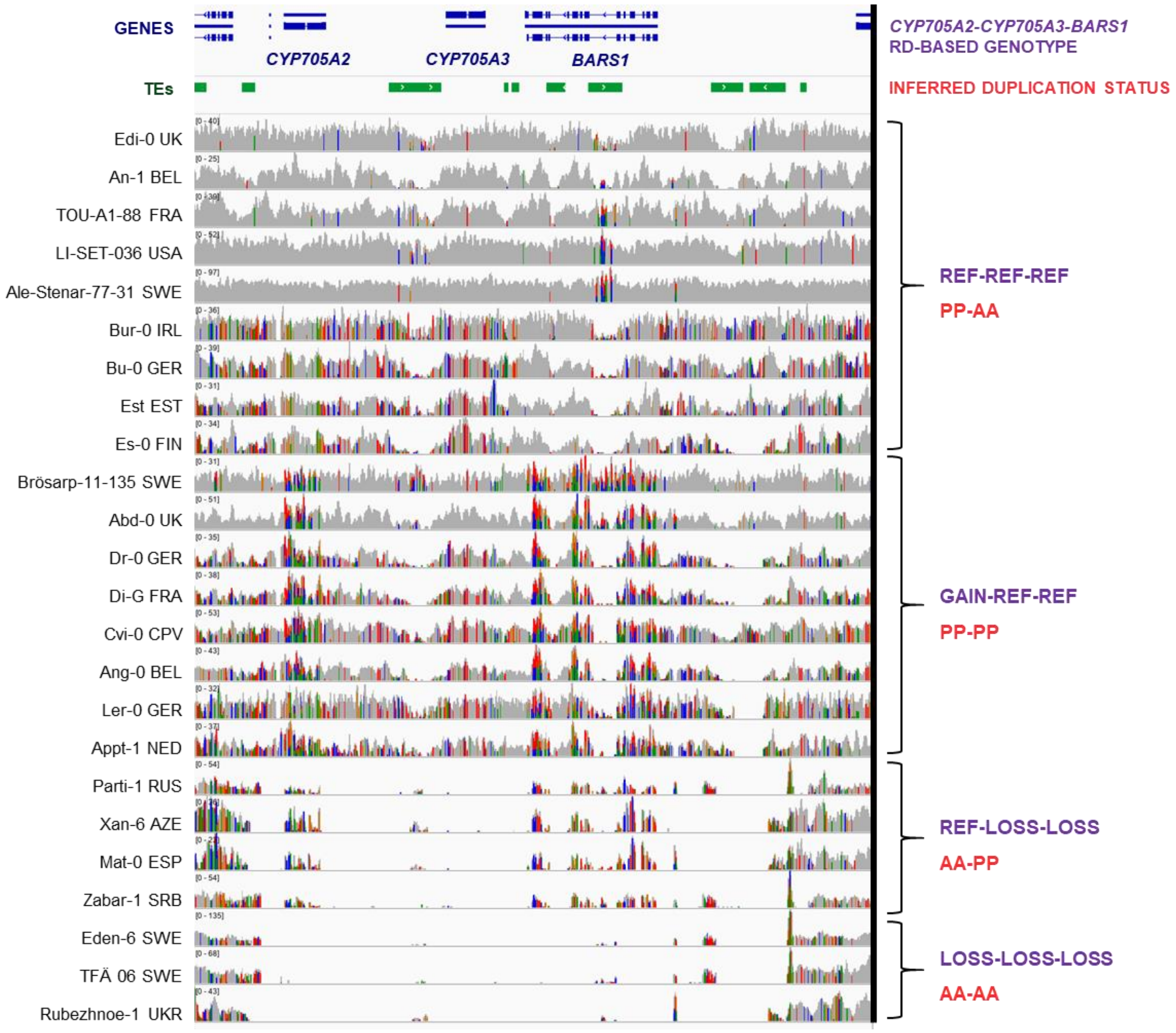


**Figure S10. Variation in WGS data coverage and mapping in the region spanning *CYP705A2, CYP705A3* and *BARS1* genes.** The presence (P) / absence (A) of *CYP705A2*, *BARS1* and their duplicates was inferred based on a combination of RD genotyping results and the SNP analysis at *CYP705A2* and *BARS1* loci (see Supplemental Table S11 and Methods for details).

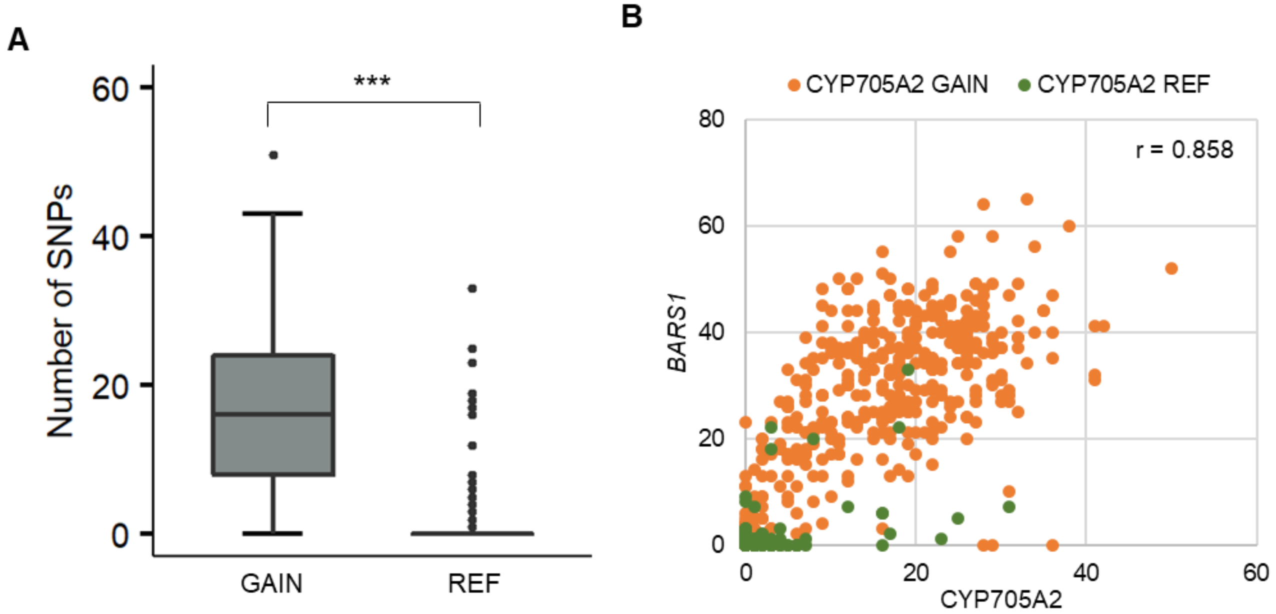


**Figure S11.** ***CYP705A2*** **duplication detected by RD assay correlates with the occurrence of heterozygous SNPs at *CYP705A2* and *BARS1* loci.** A) Number of heterozygous SNPs found in *CYP705A2* coding sequence in accessions with GAIN and REF genotypes. Boxplots show median (inner line) and inner quartiles (box). Whiskers extend to the highest and lowest values no greater than 1.5 times the inner quartile range. Asterisks indicate statistical significance (Wilcoxon rank sum test with continuity correction, ***p.value <0.001). B) Correlation of heterozygous SNP frequency between *CYP705A2* and *BARS1*. Colors indicate accessions with varying *CYP705A2* copy numbers. r – Pearson's correlation coefficient.

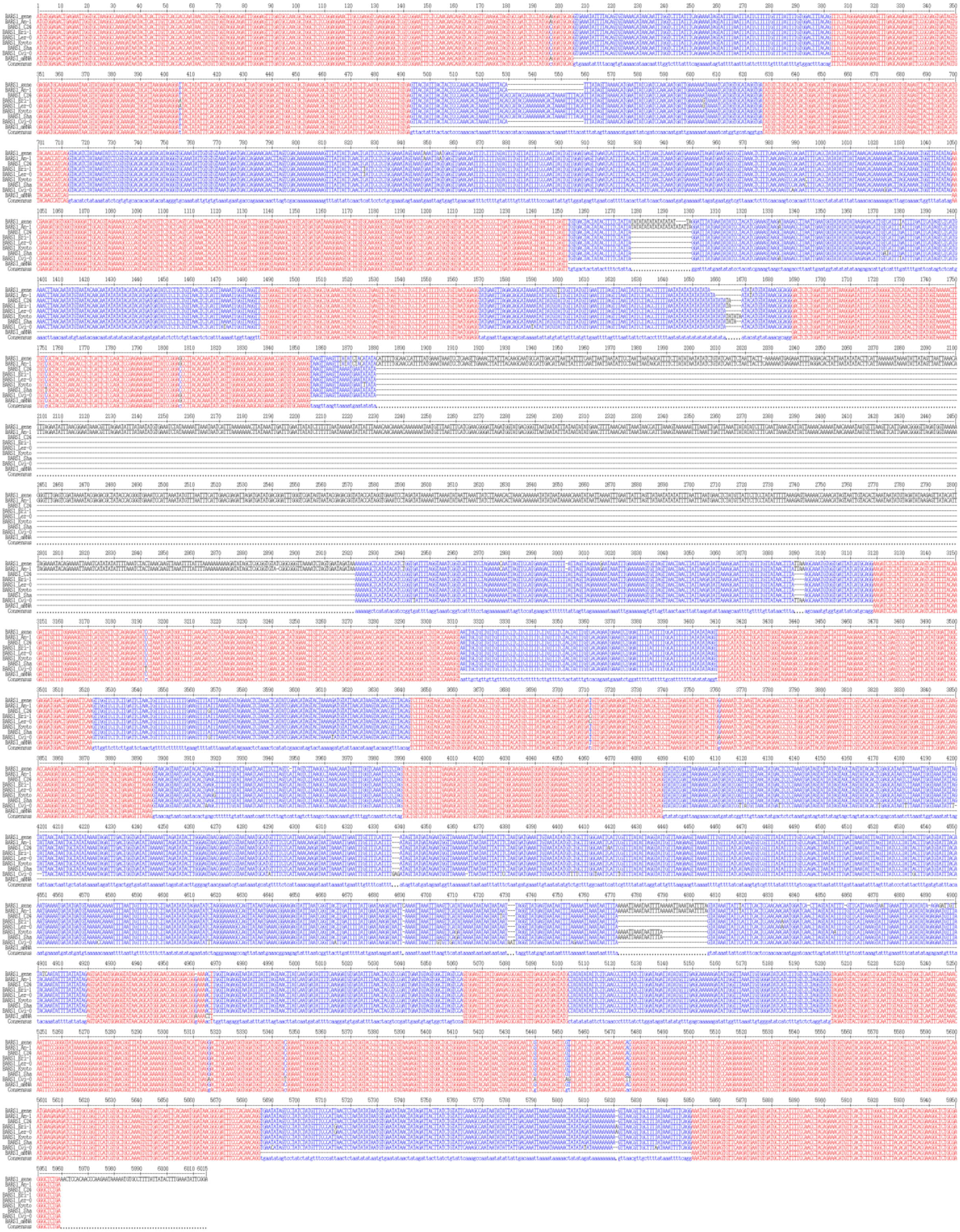

**Figure S12. Multiple sequence alignment of *BARS1* genomic sequences reveals a common lack of the largest intron.** Sequence order from the top: Col-0; An-1; C24; Eri-1; Ler-0; Kyoto; Sha; Cvi-0; BARS1_mRNA (to indicate exon-intron organization) and consensus.

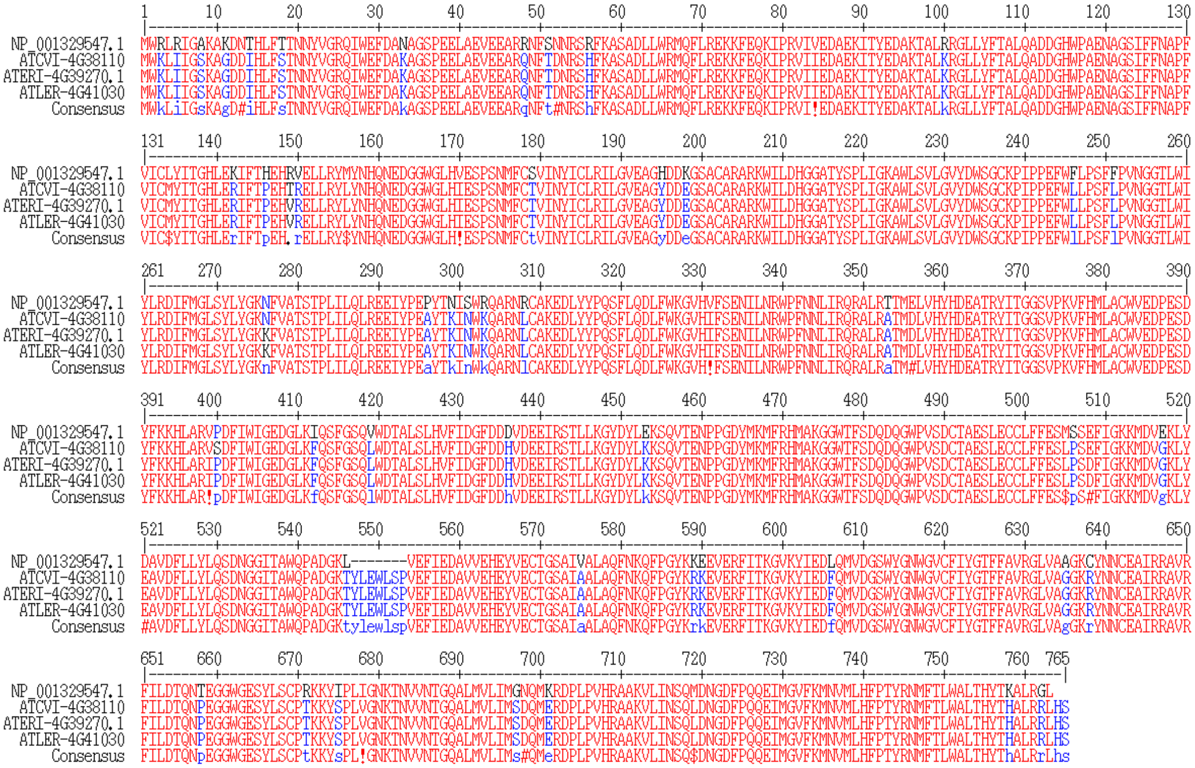


**Figure S13. Comparison of baruol synthase 1 protein NP_001329547.1 with proteins encoded by *BARS2* genes in Cvi-0, Eri-1 and Ler-0.** Multiple sequence alignment generated with Multalin with default parameters.

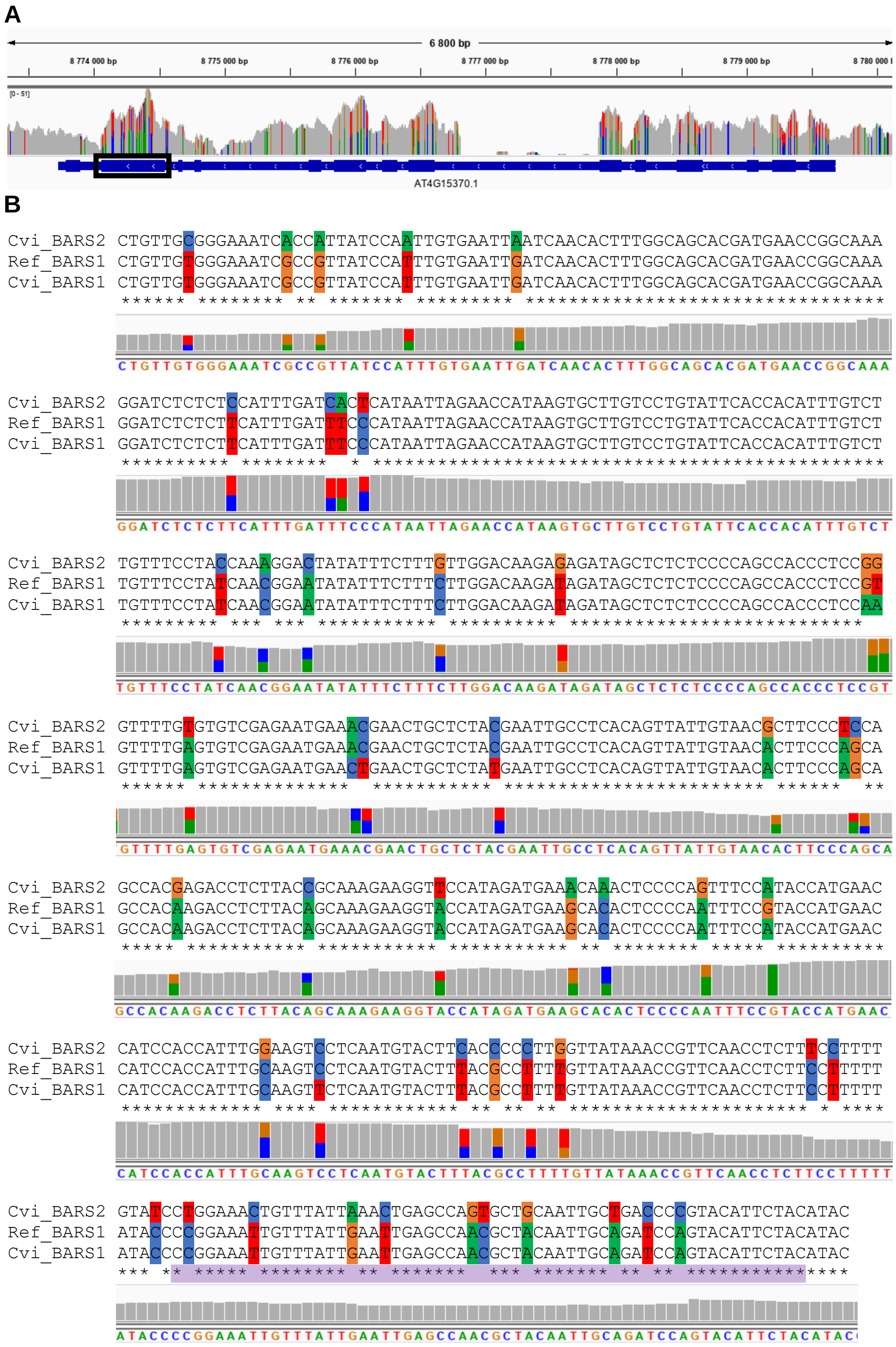


**Figure S14. Heterozygous SNPs in Cvi-0 co-localize with sequence differences between BARS1and its duplicate.** A) Read coverage at BARS1 locus for Cvi-0 (reads mapped to the reference genome). B) Multiple sequence alignment of the *BARS1* fragment (region chr4:8774060-8774546, marked by the black box in A) with *BARS1* and its duplicate (denoted as *BARS2*) from Cvi-0. IGV screenshots presenting Cvi-0 read coverage and the SNP positions are overlaid. The purple block indicates the position of the C080 MLPA probe.

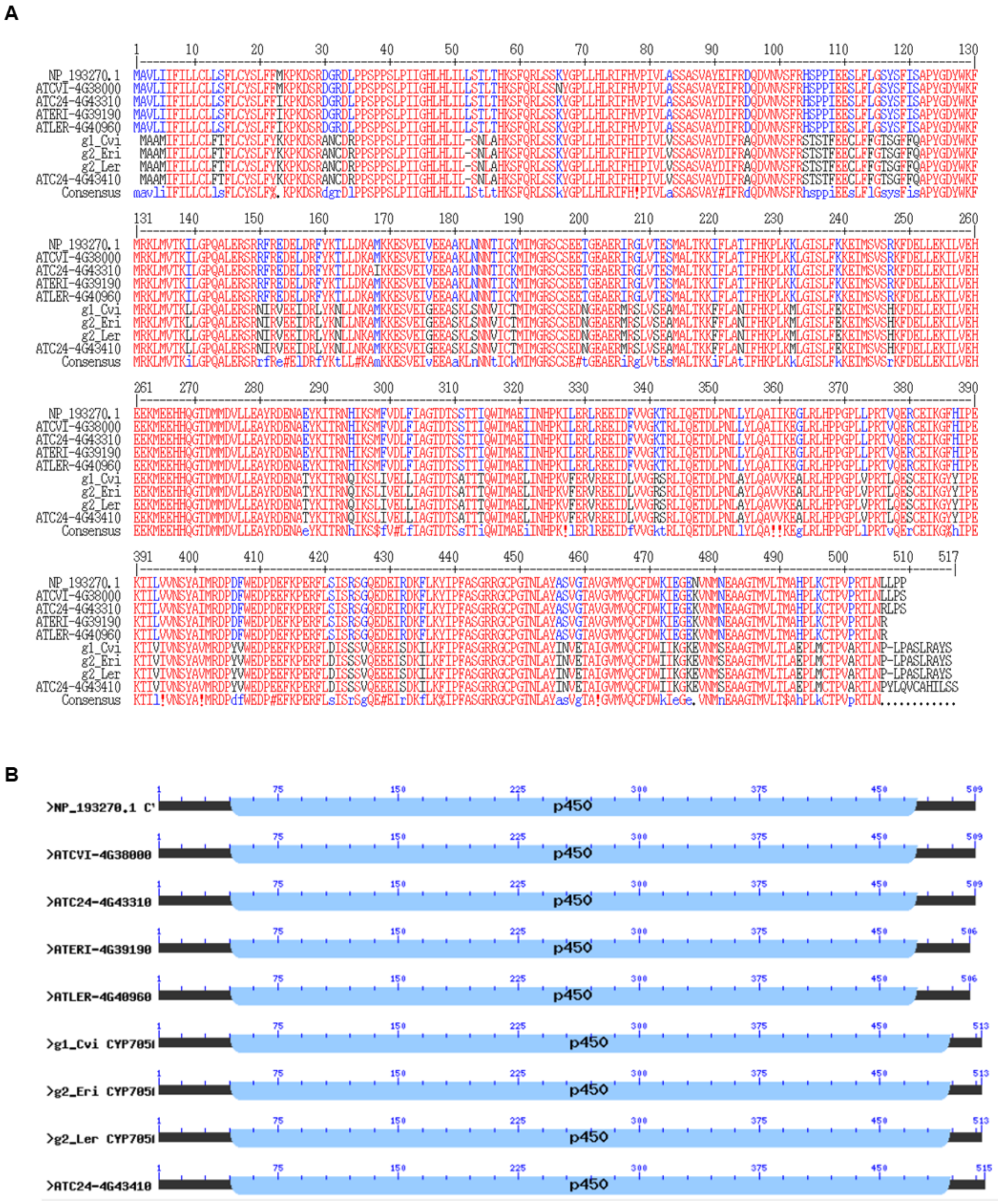

**Figure S15. Sequence comparison of *CYP705A2* and its duplicate *CYP705A2a*.** A) Multiple protein alignment of the proteins encoded by *CYP705A2* in Col-0, Cvi-0, C24, Eri-1 and Ler-0 (NP_193270.1, ATCVI-4G38000, ATC24-4G43310, ATERI-4G39190, ATLER-4G40960, respectively) and proteins encoded by CYP705A2 in Cvi-0, Eri-1, Ler-0 and C24 (g1_Cvi, g2_Eri, g2_Ler, ATC24-4G43410, respectively). B) Conserved protein domains in CYP705A2 and CYP705A2a sequences found by searching the Pfam database.

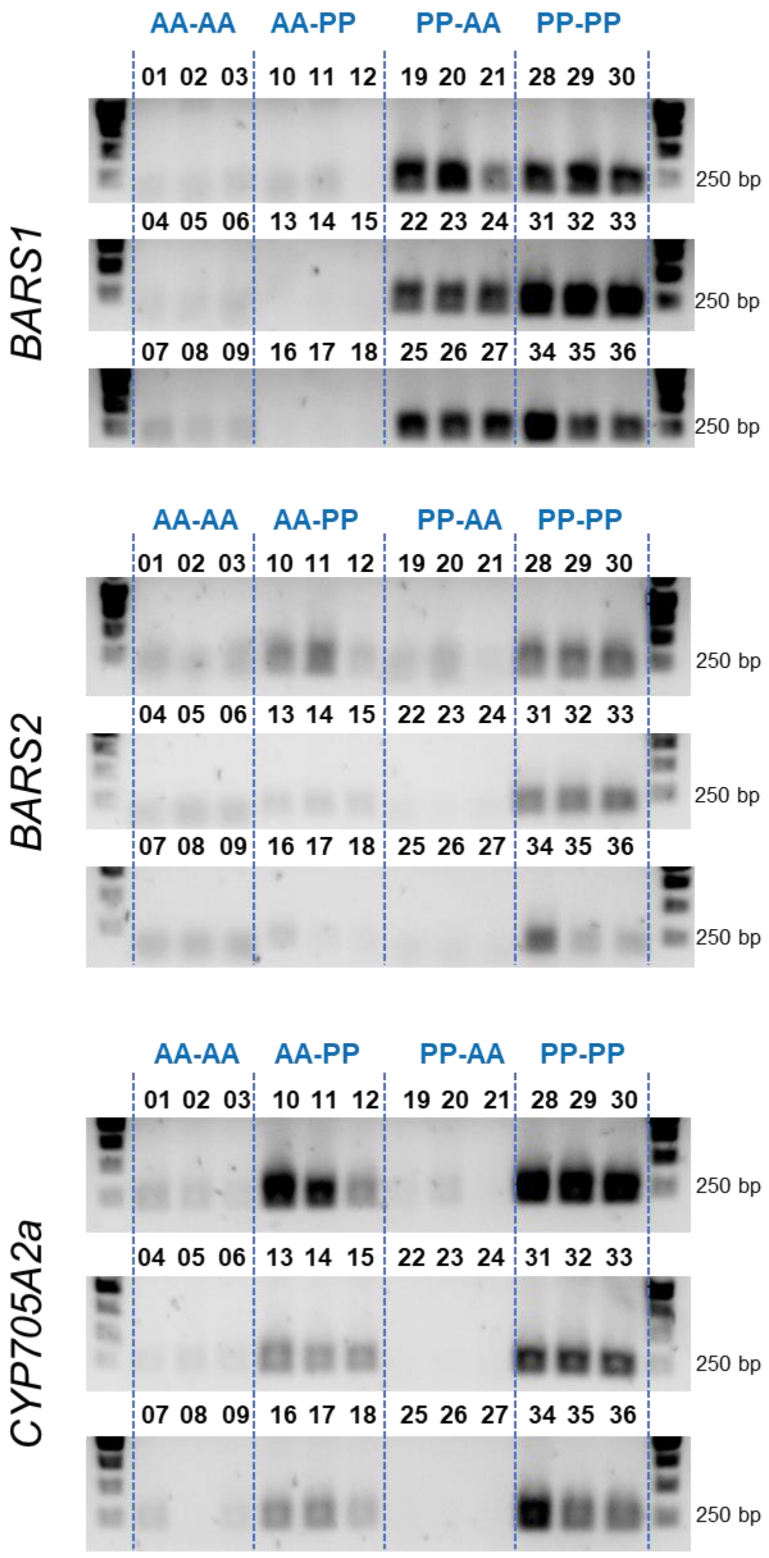


**Figure S16. PCR verification of group assignments based on the presence/absence of *BARS1, CYP705A2a* and *BARS2* genes.** Sample identities are provided in Supplemental Table S11.

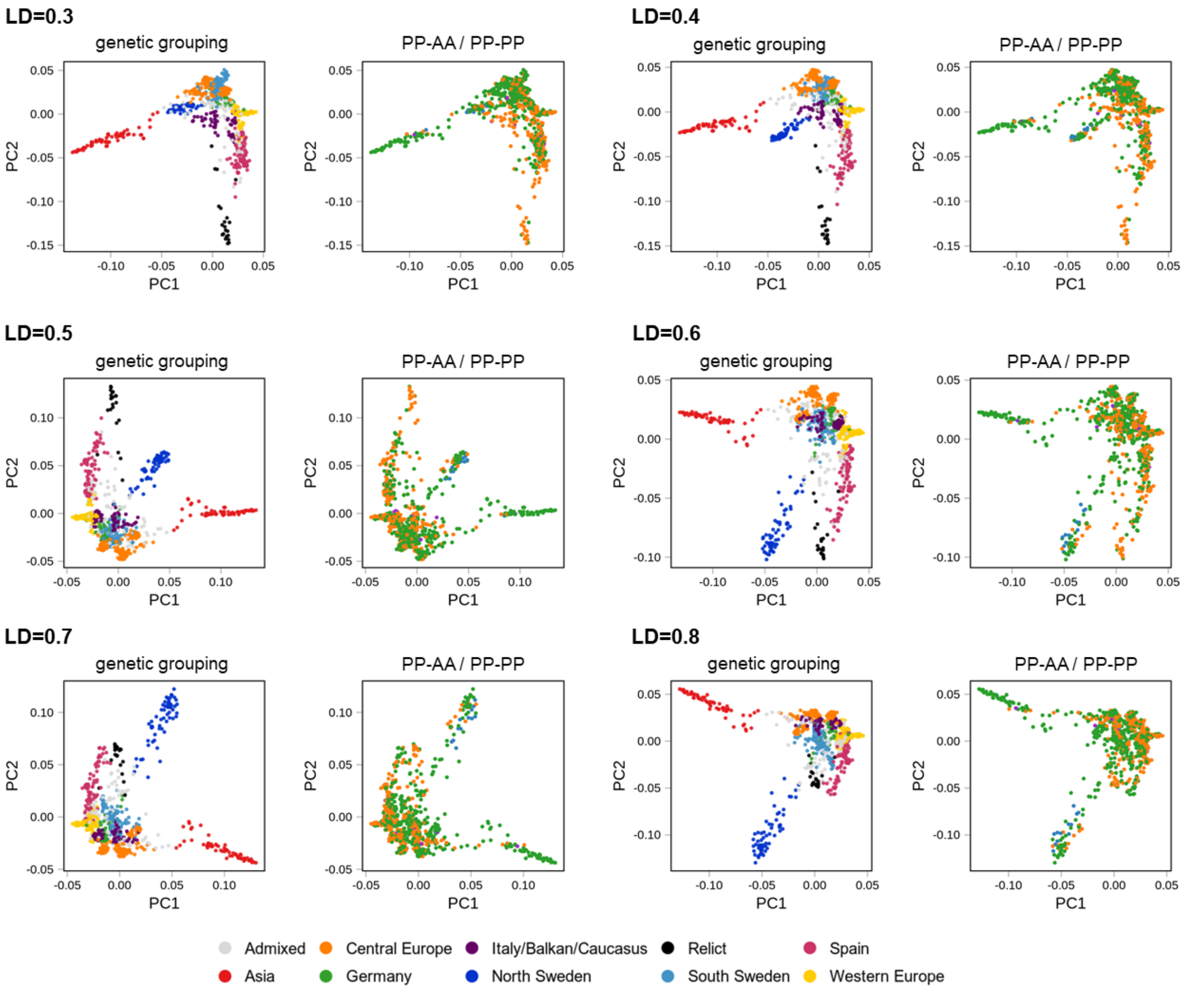


**Figure S17. Spread of PP-AA and PP-PP variants of arabidiol/baruol gene cluster in Arabidopsis population.** Principal component analysis (PCA) plots were generated with varying LD parameter. U.S.A. accessions were excluded from the analysis to better visualize other groups. Plots are colored according to main genetic groups (left) or *CYP705A2-BARS1* duplication status (right). Plots generated at LD=0.3 are also presented in Fig. 4 in the main text.

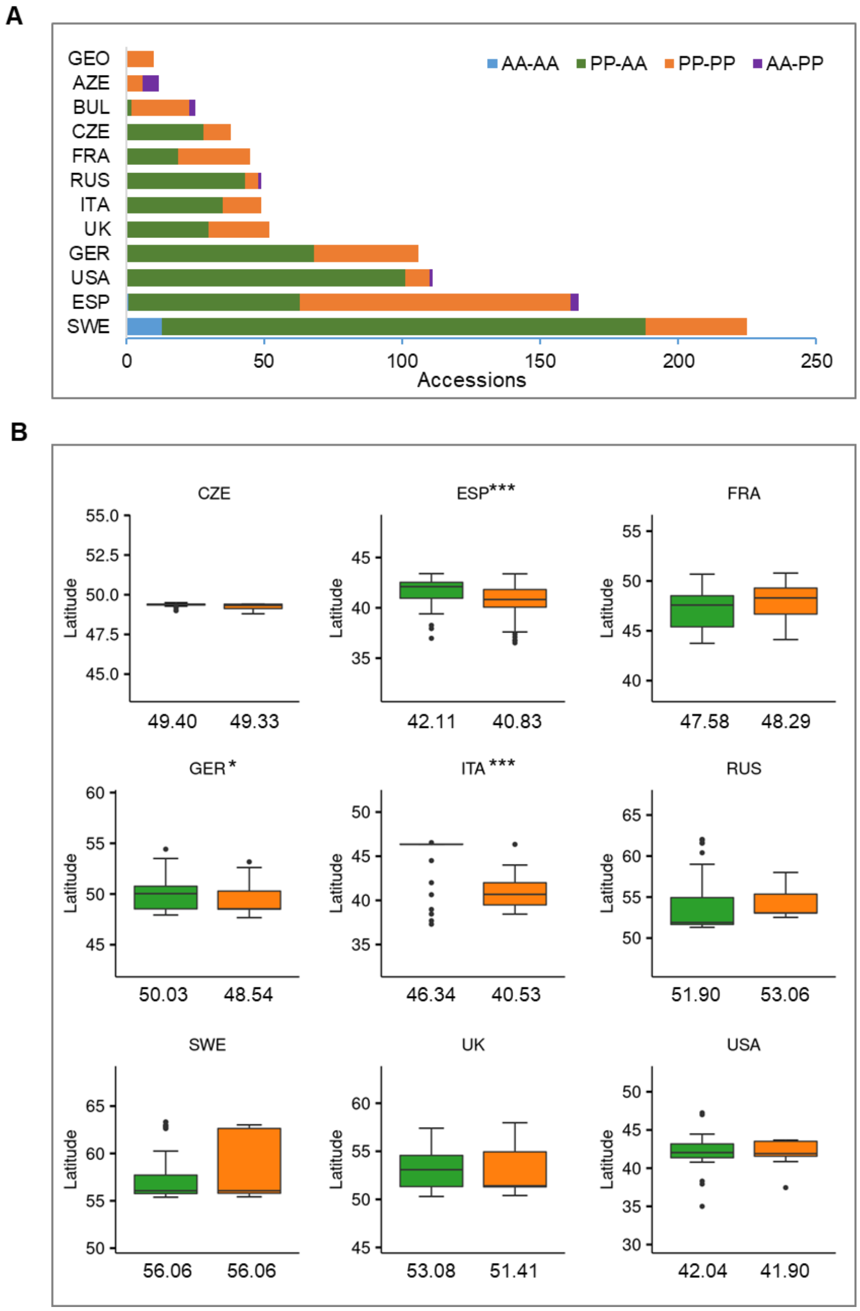


**Figure S18. Latitudes of origin among accessions with and without *CYP705A2a-BARS2* genes divided by country.** A) Frequency of four groups in individual countries. B) Boxplots presenting collection site latitudes of accessions from PP-AA and PP-PP groups. Only countries with ≥5 accessions within each group are presented. Median values are presented below the boxplots. Boxplots show median and inner quartiles. Whiskers extend to the highest and lowest values no greater than 1.5 times the inner quartile range. Asterisks indicate statistical significance (Wilcoxon rank sum test with continuity correction, *p.value<0.05, ***p.value <0.001).

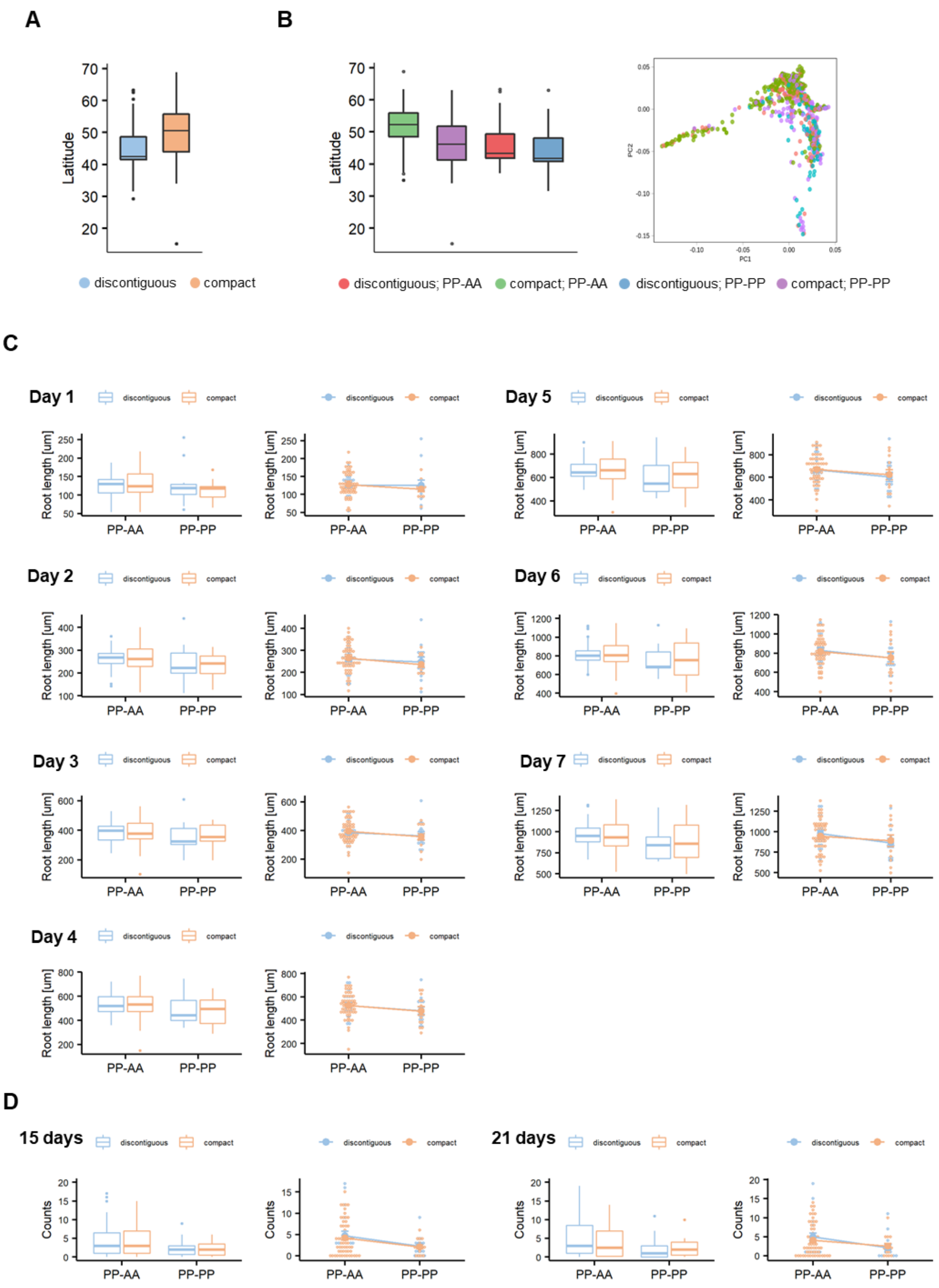


**Figure S19. The combined effect of thalianol and arabidiol/baruol gene clusters' structural variation on root growth phenotypic variation.** A) Association of variability in thalianol gene cluster organization with latitude B) PCA on >200k SNPs and LD = 0.3 (left) and latitudes of origin (right) of accessions divided by both thalianol and arabidiol/baruol cluster type. C,D) Root growth phenotypes presented in the main text in Fig. 5B and 5C, respectively, divided by both thalianol and arabidiol/baruol cluster type (left), along with two-way ANOVA plot (right).

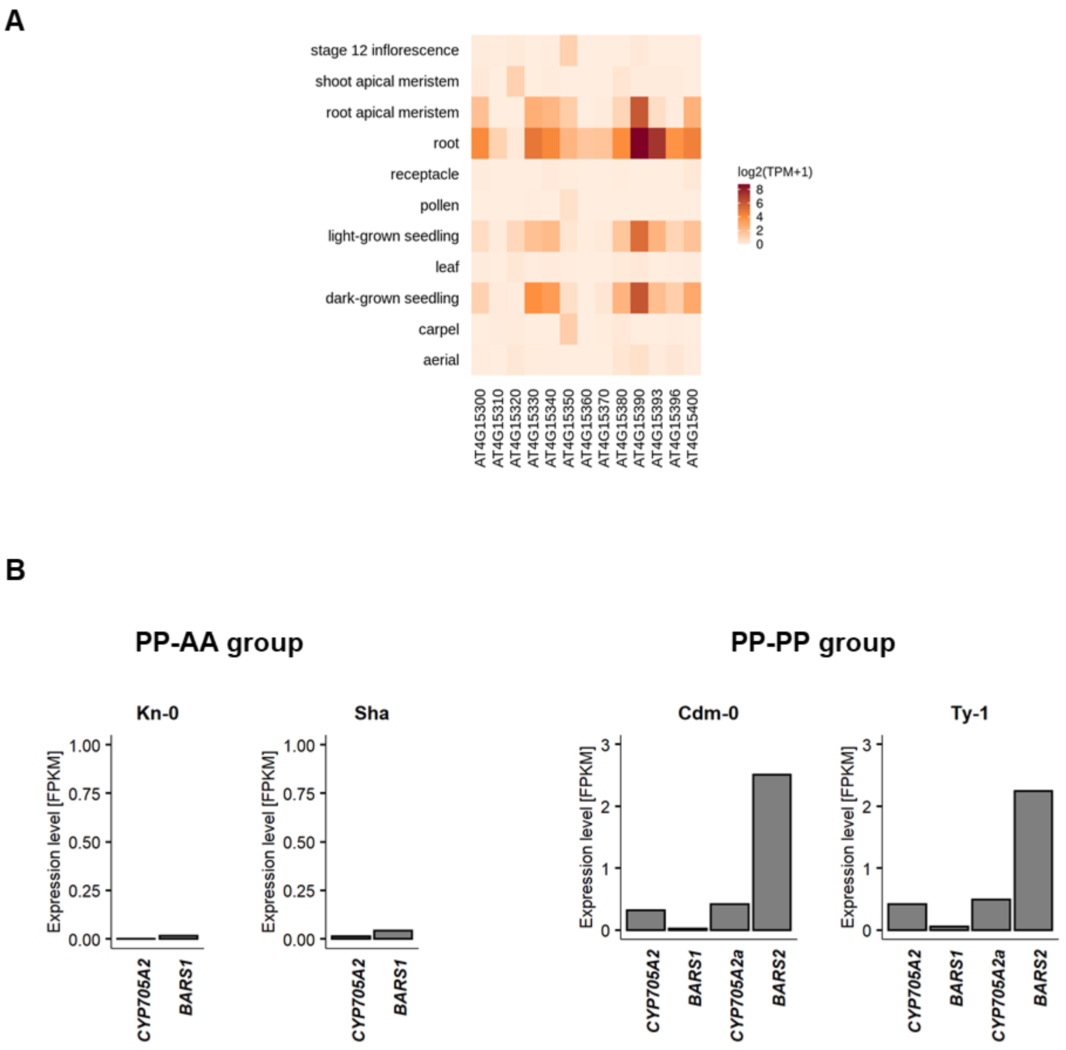


**Figure S20. Differences in expression of arabidiol/baruol gene cluster in among accessions.** A) Tissue-specific expression in Col-0 accession (PP-AA group). B) Expression of *CYP705A2*, *BARS1*, *CYP705A2a* and *BARS2* genes in accessions from PP-AA and PP-PP groups. For each accession, RNA-Seq data were mapped to the respective genomic assemblies. Where necessary, the region of interest was annotated with Augustus and the gene models were used for FPKM calculations.

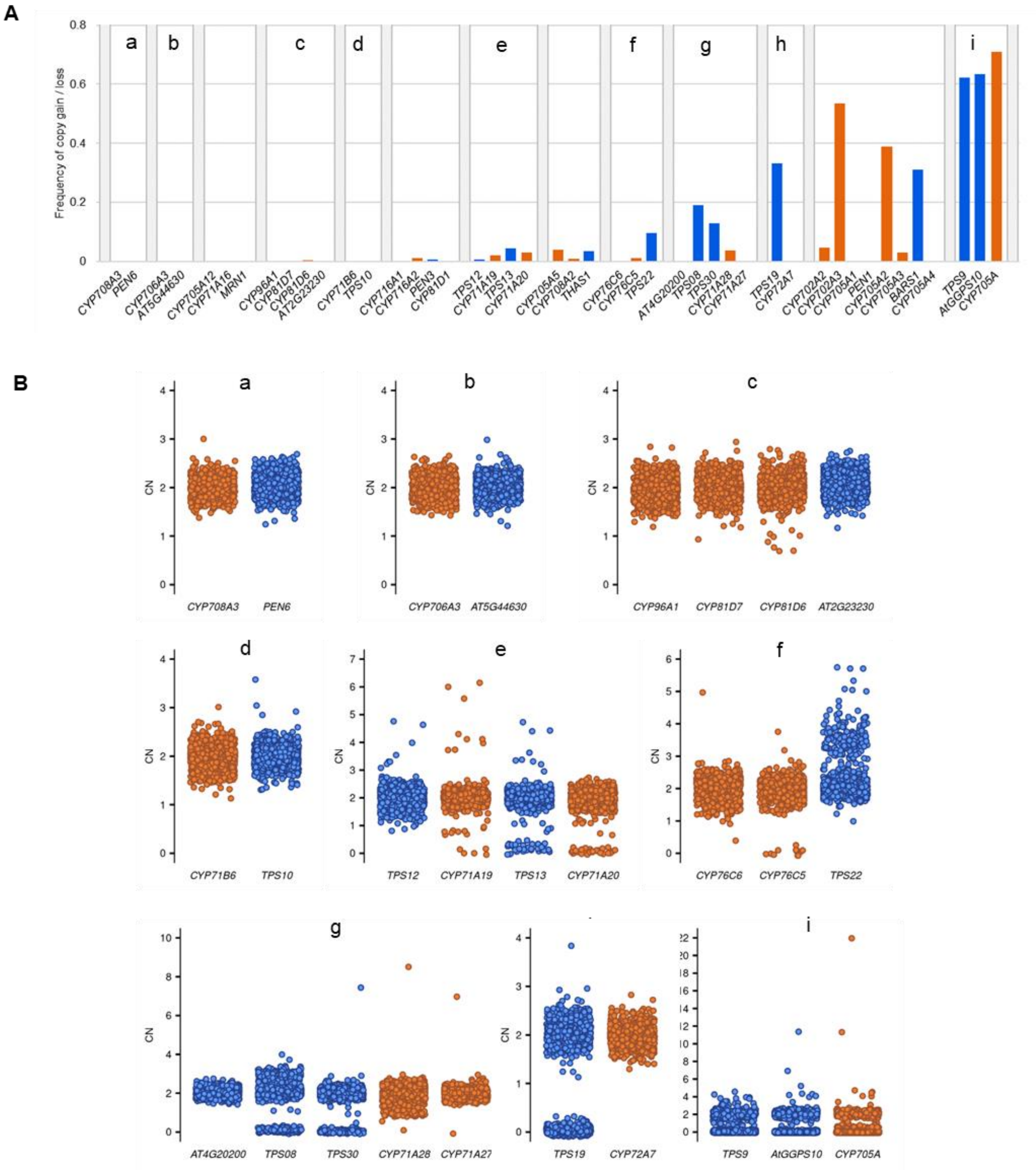


**Figure S21. Copy number variation TS/CYP gene pairs in Arabidopsis genome.** A) Frequency of gene copy number changes. All TSs and CYPs that form gene pairs are shown; B) Distribution of RD-based genotypes among the accessions. Pairs were defined using a 30-kb distance cutoff. Pairs sharing a CYP or TS gene were grouped together. Pairs from thalianol, marneral, tirucalladienol and arabidiol/baruol gene clusters are not displayed.
